# A Distinct Layer 1 Astrocyte Program Shapes Perisynaptic Structure and Calcium Signaling in Mouse Motor Cortex

**DOI:** 10.64898/2026.03.16.712002

**Authors:** Srimayee Bhattacharjee, Kun-Hai Yeh, Ping-Yen Wu, Ming-Xi Pan, Tzu-Hsien Liu, Zhao-Bo Tsai, Sok-Keng Tong, Zi-Hui Zhuang, Yu-Min Huang, Shen-Ju Chou, Shu-Ling Chiu, Ming-Yi Chou, Chen-Hsin Albert Yu, Yu-Wei Wu

## Abstract

Although layer-specific molecular and morphological diversity among cortical astrocytes is increasingly well established, how these distinct states are linked to specialized calcium signaling and maintained in the adult cortex remains unclear. Here, combining super-resolution structural imaging, two-photon calcium imaging, and analysis of public single-cell and spatial transcriptomic resources, we identify Layer 1 (L1) astrocytes in mouse primary motor cortex as a distinct superficial astrocyte program. These cells occupy compact territories yet contain dense synapse-associated loop-like structures and display frequent, fast, broadly spreading calcium events that engage a large fraction of the territory. Transcriptomic analyses identify *Id1* and *Id3* as enriched components of this superficial program. CRISPR–Cas9 deletion of *Id1* and *Id3* in adult astrocytes selectively disrupts L1 and superficial Layer 2/3 (L2/3) astrocytes, expanding territory size, reducing fine-process complexity, and suppressing calcium activity. Thus, adult layer-specific transcriptional programs maintain specialized astrocyte structure and signaling.

**Graphical Abstract:** 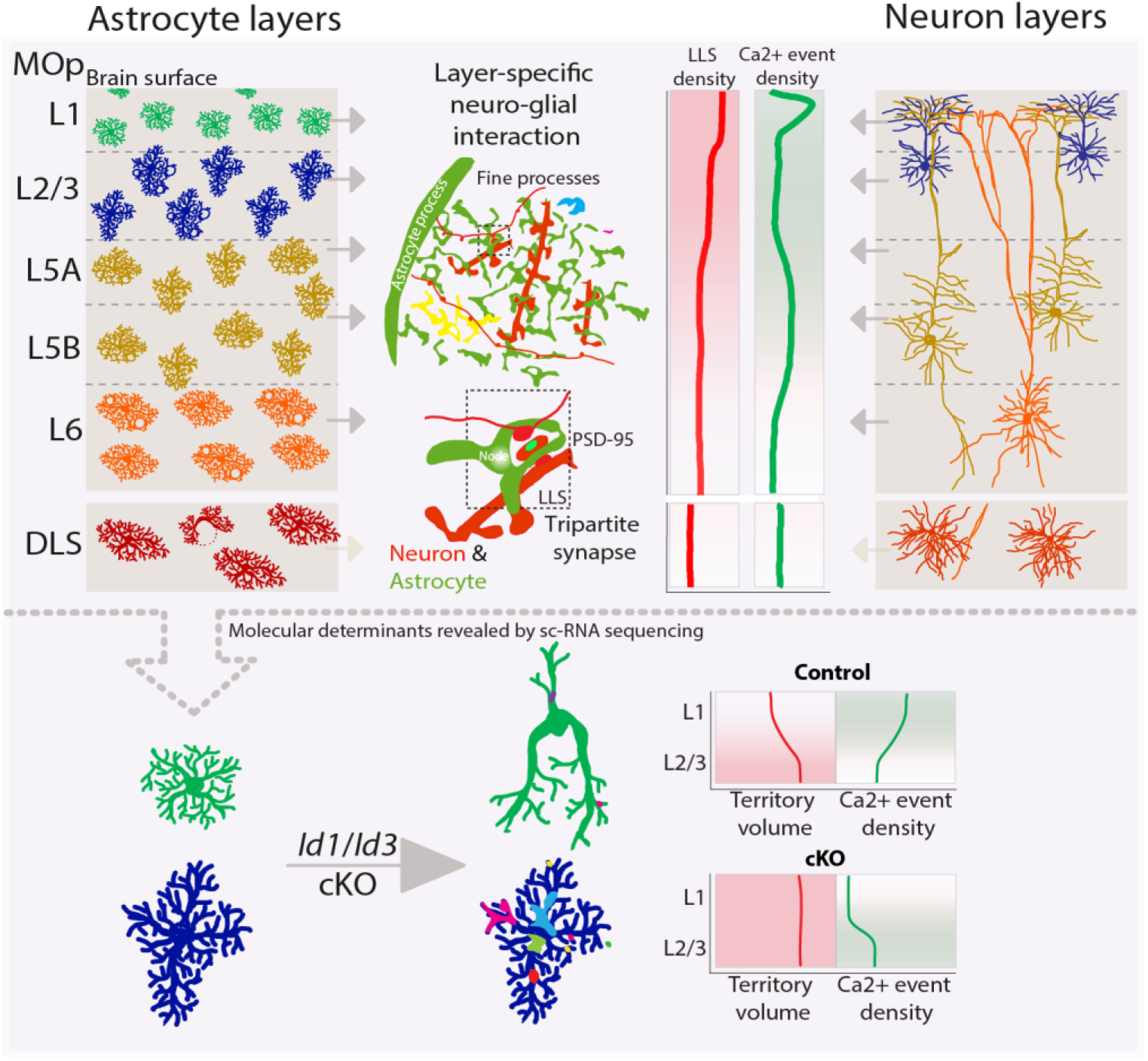

**Highlights:**

- Layer 1 astrocytes in mouse motor cortex have compact, loop-rich nanoarchitecture
- Layer 1 astrocytes show fast, frequent, widespread calcium events
- Single-cell data define a superficial astrocyte state enriched for *Id1* and *Id3*
- Adult *Id1/Id3* loss selectively disrupts Layer 1 astrocyte structure and calcium

**In Brief:** Bhattacharjee *et al.*, identify a specialized Layer 1 astrocyte state in mouse motor cortex, defined by compact, loop-rich nanoarchitecture and unusually high calcium signaling. *Id1*/*Id3* deletion selectively disrupts this superficial program, linking layer-specific astrocyte form and function to adult transcriptional control.

## Introduction

Layer-specific diversity of cortical astrocytes is now well established. Single-cell and spatial transcriptomic studies, together with anatomical analyses, have identified molecularly and morphologically distinct astrocyte states across cortical depth and across brain regions^1–11^. This view challenges the long-standing notion of astrocytes as a uniform support population and instead suggests that they are adapted to the specific demands of local circuits. Such specialization is likely to be functionally important because astrocytes regulate synapse formation, neurotransmitter recycling, and cerebral blood flow^12–15^. Yet, for most cortical astrocyte states, how molecular identity and structural specialization translate into distinct signaling properties remains unresolved.

The primary motor cortex (MOp) provides a particularly powerful setting in which to study layer-specific astrocyte specialization. Multimodal anatomical and spatial transcriptomic atlases show that MOp contains refined sublaminar organization, depth-dependent cell populations, and distinct input-output architecture across cortical depth, creating markedly different local circuit contexts across layers^1,16,17^. Astrocytes in the MOp are also functionally engaged in motor behavior: attenuating astrocytic calcium signaling impairs motor-skill learning and MOp synaptic plasticity, astrocyte-specific manipulation of GLT1 or Gq signaling disrupts learned movement execution and neuronal ensemble coding, and recent *in vivo* imaging shows that astrocytic calcium dynamics in MOp are modulated during motor learning^18–20^. More broadly, cortical astrocytes can form behavior-dependent calcium hotspot maps with day-to-day stability, consistent with persistent subcellular specialization^21^. Together, these observations suggest that MOp astrocytes are embedded in layer- and behavior-specific circuit operations, but how layer-specific astrocyte states are coupled to fine-process nanoarchitecture and calcium signaling remains unclear.

A central readout of astrocyte specialization is intracellular calcium signaling. Although astrocytes are electrically non-excitable, they display diverse calcium dynamics in response to synaptic and neuromodulatory activity^22–28^, and perturbing these signals alters circuit function and behavior^29–33^. Prior work has also shown that astrocyte calcium dynamics differ across cortical layers and across brain regions^11,34–36^, indicating that astrocytes are not functionally interchangeable. What remains less clear is how these distinct signaling modes relate to the nanoscale organization of astrocytic processes. Fine loop-like structures (LLSs) within astrocytic arbors are closely associated with synapses and can serve as hotspots for local calcium activity^37,38^, raising the possibility that layer-specific nanoarchitecture helps determine how astrocytes sample and integrate local synaptic inputs.

Layer-specific astrocyte states are likely shaped not only by local circuit environments^39,40^ but also by intrinsic programs that stabilize them in adulthood. Neuronal layering influences cortical astrocyte gene expression^3,4^, and region-specific transcriptional control is increasingly recognized as an important determinant of astrocyte morphology and function^41,42^. Among candidate regulators, the *Id* family members *Id1* and *Id3* are well known for their roles in the gliogenic switch and in balancing proliferation and differentiation^43–46^. Prior transcriptomic studies have further suggested superficial enrichment of these genes in cortical astrocytes^3^, pointing to a possible role in the maintenance of specialized astrocyte states after development. Whether *Id1* and *Id3* actively sustain layer-specific astrocyte structure and signaling in the adult cortex, however, remains unknown.

Here, we combine super-resolution structural imaging, two-photon calcium imaging, and analysis of publicly available single-cell and spatial transcriptomic datasets to define layer-specific astrocyte specialization in adult mouse MOp. We identify L1 astrocytes as a distinct superficial astrocyte program characterized by compact territories, dense synapse-associated LLS nanoarchitecture, and unusually frequent, fast, and broadly spreading calcium events. Transcriptomic analyses link this phenotype to an L1-enriched program featuring *Id1* and *Id3*, and CRISPR-Cas9-mediated deletion of these genes in adult astrocytes selectively disrupts superficial astrocyte morphology and calcium activity. Together, these findings connect layer-specific astrocyte identity to specialized structure, signaling, and adult transcriptional maintenance.

## Results

### Heterogeneity in astrocytic morphology across layers of the MOp and the DLS

To ask whether astrocytes are organized into distinct spatial states within the motor circuit, we first performed a layer-resolved morphological survey across the mouse MOp and the dorsolateral striatum (DLS). In tamoxifen-induced Aldh1l1-CreER^T2^;LSL-tdTomato mice, sparse astrocyte labeling enabled two-photon imaging and three-dimensional reconstruction of individual cells from L1, L2/3, Layer 5 (L5), Layer 6 (L6), and DLS in acute coronal slices (**Figures 1A–1D** and **S1**). Laminar identity was assigned using IR-differential interference contrast (IR-DIC) landmarks (**Figure S1**), and we quantified projected territory area, territory volume, and arbor orientation relative to the brain surface (**Figures 1E–1G**).

**Figure 1.**
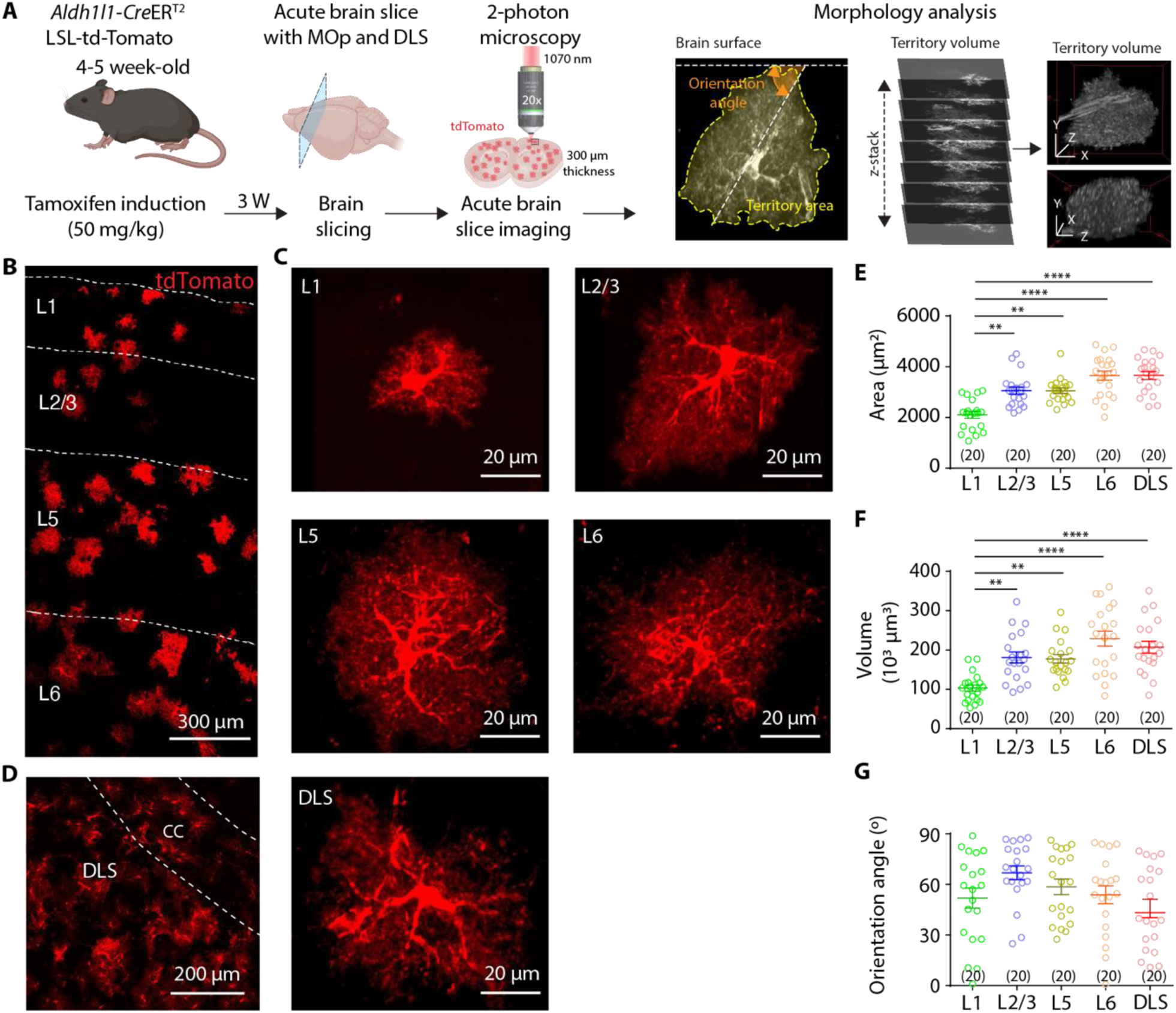
Detailed morphology study of astrocytes in the MOp and DLS. (A) Experimental outline highlighting the steps from tamoxifen-induction to imaging analysis. (B) Sparse labeling of astrocytes expressing tdTomato in the L1, L2/3, L5 and L6 of the MOp. Scale bar represent 300 µm. (C) Representative examples of astrocytes imaged from these regions. Scale bars represent 20 µm. (D) Left: sparse labeling of astrocytes expressing tdTomato in the Corpus Callosum (CC) and the DLS. Scale bar represent 300 µm. Right: representative example of astrocyte imaged from the DLS. Scale bar represent 20 µm. (E) Plot showing the cross-section area of the imaging plane occupied by single astrocyte territory across the MOp and DLS (L1: 2106 ± 137 µm^2^, n = 20; L2/3: 3056 ± 145 µm^2^, n = 20; L5: 3054 ± 107 µm^2^, n = 20; L6: 3655 ± 183 µm^2^, n = 20; DLS: 3659 ± 155 µm^2^, n = 20 cells; N = 14 mice; Kruskal-Wallis test followed by Dunn’s multiple comparisons test). (F) Plot showing the volume occupied by single astrocyte territory across the MOp and DLS (L1: 103 ± 8 (x10^3^ µm^3^), n = 20; L2/3: 181 ± 14 (x10^3^ µm^3^), n = 20; L5: 178 ± 11 (x10^3^ µm^3^), n = 20; L6: 229 ± 19 (x10^3^ µm^3^), n = 20; DLS: 207 ± 15 (x10^3^ µm^3^), n = 20 cells; N = 14 mice; Kruskal-Wallis test followed by Dunn’s multiple comparisons test). (G) Plot showing the orientation angle of astrocytes with respect to the brain surface (L1: 52 ± 6°, n = 20; L2/3: 67 ± 4 °, n = 20; L5: 59 ± 5 °, n = 20; L6: 54 ± 5 °, n = 20; DLS: 46 ± 5 °, n = 20 cells; N = 14 mice; Kruskal-Wallis test followed by Dunn’s multiple comparisons test). The data are shown as mean ± SEM. **p < 0.01, ****p < 0.0001.

This analysis immediately distinguished the superficial cortex from the rest of the motor circuit. Among all regions examined, L1 astrocytes occupied the smallest territories by both projected area and volume—roughly half the volume of L6 astrocytes—whereas deep-layer and DLS astrocytes occupied the largest domains (**Figures 1E** and **1F**). Thus, astrocyte morphology did not vary as a simple continuum across depth; instead, L1 emerged as a clear anatomical outlier. By contrast, orientation relative to the brain surface showed only modest, non-significant variation across regions (**Figure 1G**), indicating that laminar specialization is reflected more strongly in territorial scale than in overall arbor alignment. Because L1 is largely devoid of neuronal somata and instead contains apical tuft dendrites and synapse-rich neuropil, whereas deeper layers and DLS contain more somatic and fiber-rich elements, these data pointed to a distinct structural niche for superficial astrocytes. This initial anatomical survey therefore identified L1 astrocytes as candidates for a specialized superficial program and motivated us to ask whether their fine perisynaptic architecture is also uniquely organized.

### Superficial astrocytes exhibit a dense synapse-associated fine-process nanoarchitecture

The compact territorial footprint of L1 astrocytes raised the possibility that superficial cells are not simply smaller versions of deeper astrocytes, but instead organize their distal arbor differently. Because fine processes account for most astrocyte volume^47^, we used super-resolution structured illumination microscopy (SR-SIM) to examine this compartment across MOp layers and DLS. Three standardized ROIs were sampled from the fine processes of each cell (**Figures 2A** and **2B**), where we resolved loop-like structures (LLSs), closed or partially closed membrane elements within distal astrocytic arbors (**Figures 2B–2D** and **S2A**). Although LLSs were present in all regions examined, they were visually enriched in L1 astrocytes.

**Figure 2.**
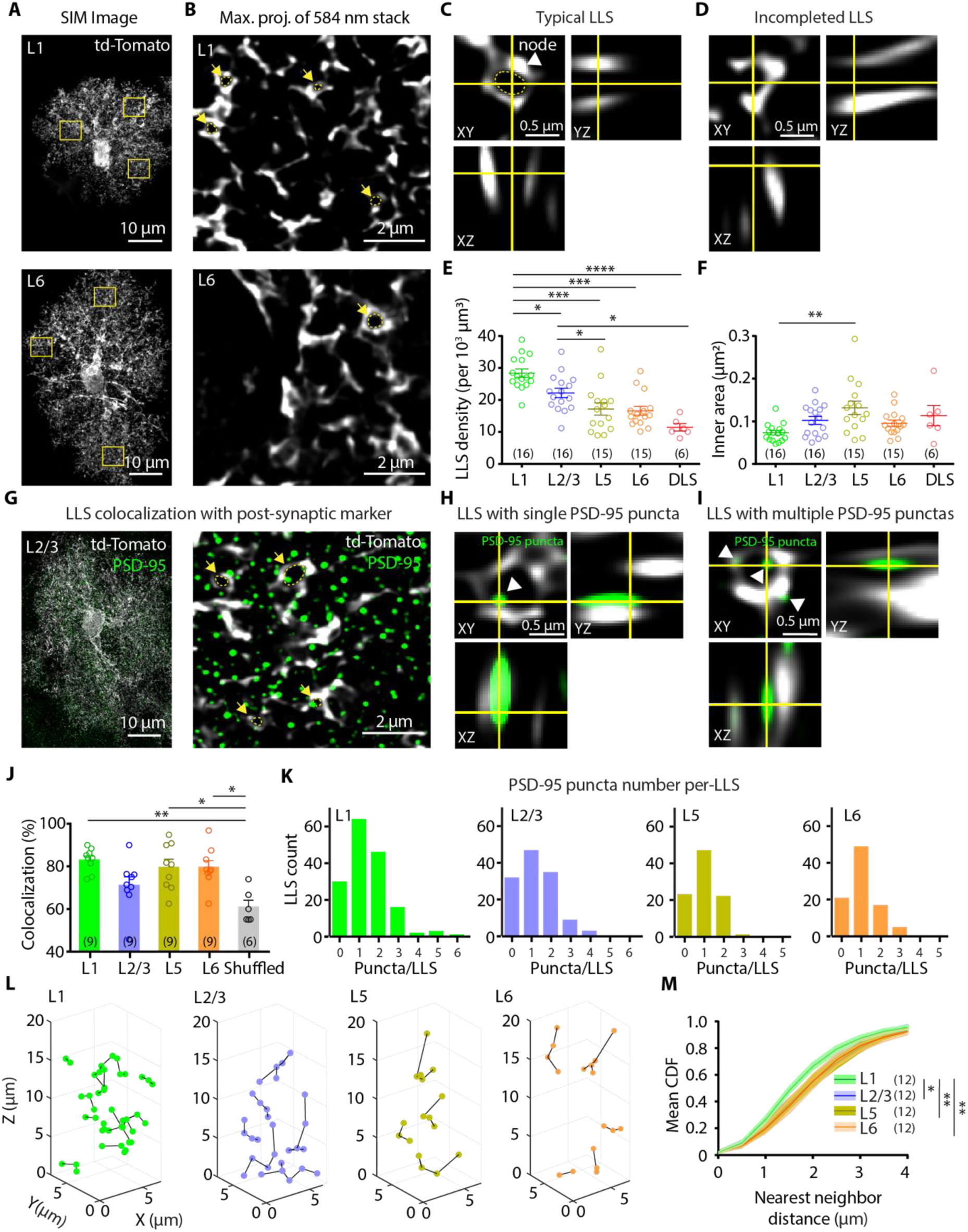
Quantification of LLS density across the MOp. (A) Top: 3D projection of a L1 astrocyte imaged using SR-SIM. Bottom: 3D projection of a L6 astrocyte imaged using SR-SIM. The yellow rectangles denote the ROIs chosen from the fine processes. Scale bars represent 10 μm. (B) Chosen ROIs displaying the loop-like structures (LLS) of L1 and L6 astrocytes (Top and bottom, respectively), highlighted in yellow, marked by yellow arrows. Scale bars represent 2 μm. (C) Orthogonal views of a typical LLS, yellow circle denoting its inner area, perimeter and the arrowhead points at the node structure. Scale bars represent 0.5 μm. (D) Orthogonal views of an incomplete LLS. Scale bars represent 0.5 μm. (E) Plot shows the LLS density across MOp and DLS. (L1: 28.4 ± 1.3 per 10^3^ µm^3^, n = 16 ROIs/ 6 cells; L2/3: 22.2 ± 1.4 per 10^3^ µm^3^, n = 16 ROIs/ 6 cells; L5: 17.2 ± 1.9 per 10^3^ µm^3^, n = 15 ROIs/ 5 cells; L6: 16.6 ± 1.4 per 10^3^ µm^3^, n = 15 ROIs/ 5 cells; DLS: 11.5 ± 1.2 per 10^3^ µm^3^, n = 6 ROIs/ 2 cells; N = 6 mice; Kruskal-Wallis test followed by Dunn’s multiple comparisons test). (F) Quantification of the inner area of these LLS. (L1: 0.073 ± 0.005 µm^2^, n = 16 ROIs/ 6 cells, L2/3: 0.102 ± 0.009 µm^2^, n = 16 ROIs/ 6 cells, L5: 0.132 ± 0.015 µm^2^, n = 15 ROIs/ 5 cells; L6: 0.095 ± 0.007 µm^2^, n = 15 ROIs/ 5 cells; DLS: 0.113 ± 0.024 µm^2^, n = 6 ROIs/ 2 cells; N = 6 mice; Kruskal-Wallis test followed by Dunn’s multiple comparisons test). (G) Left: 3D projection of a L2/3 astrocyte stained with PSD-95, imaged using SR-SIM. Scale bars represent 10 μm. ROI showing merged image of LLS and PSD-95, the yellow arrows denote the LLSs co-localizing with PSD-95 puncta (right). Scale bars represent 2 μm. (H) Positive co-localization of LLS with PSD-95 puncta and their orthogonal views. Scale bars represent 0.5 μm. The green dots here represent PSD-95 staining. (I) LLS encompassing multiple PSD-95 puncta. Scale bars represent 0.5 μm. (J) Comparison of LLS-PSD-95 co-localization across the MOp with the random shuffled data, the solid bars represent the colocalization with PSD-95 (L1: 83 ± 2 %, n = 9 ROIs/ 3 cells; L2/3: 71 ± 4%, n = 9 ROIs/ 3 cells; L5: 80 ± 4 %, n = 9 ROIs/ 3 cells; L6: 80 ± 3 %, n = 9 ROIs/ 3 cells, Shuffled: 61 ± 3%, n = 3 ROIs/ 1 cell; N = 4 mice; Kruskal-Wallis test followed by Dunn’s multiple comparisons test). (K) Frequency distribution of loops and their co-localization with none to multiple PSD-95 puncta count. (L) Three-dimensional visualization of the LLSs within a ROI, shown across regions. Each sphere represents a single LLS and the black lines connect the nearest LLSs to each other. (M) Quantification of the nearest neighbor distances of LLSs across regions. (n = 4 astrocytes; 12 ROIs (L1), n = 4 astrocytes; 12 ROIs (L2/3), n = 4 astrocytes; 12 ROIs (L5), n = 4 astrocytes; 12 ROIs (L6; N = 3 mice; Kolmogorov-Smirnov test). The data are shown as mean ± SEM. *p < 0.05, **p < 0.01, ***p < 0.001, ****p < 0.0001.

Quantitative analysis revealed a striking superficial-to-deep gradient in LLS organization. L1 astrocytes displayed the highest LLS density (28.4 ± 1.3 LLSs per 10^3^ µm^3^), which declined through L2/3, L5, and L6 and reached a minimum in DLS (11.5 ± 1.2 LLSs per 10^3^ µm^3^; **Figure 2E**). Thus, the superficial compartment is distinguished not only by reduced territorial volume (**Figure 1**), but also by a disproportionately dense fine-process nanoarchitecture within that restricted space. Moreover, L1 LLSs exhibited the smallest inner areas and perimeters (**Figures 2F**; L1: 0.073 ± 0.005 µm^2^, n = 16 ROIs, L2/3: 0.1021 ± 0.009 µm^2^, n = 16 ROIs, L5: 0.1316 ± 0.015 µm^2^, n = 15 ROIs, L6: 0.0954 ± 0.007 µm^2^, n = 15 ROIs, DLS: 0.1133 ± 0.023 µm^2^, n = 6 ROIs; N = 6 mice, Kruskal-Wallis test followed by Dunn’s multiple comparisons test and **S2B;** L1: 0.9902 ± 0.030 µm, n = 16 ROIs, L2/3: 1.157 ± 0.057 µm, n = 16 ROIs, L5: 1.331 ± 0.092 µm, n = 15 ROIs, L6: 1.135 ± 0.051 µm, n = 15 ROIs, DLS: 1.309 ± 0.173 µm, n = 6 ROIs; N = 6 mice, Kruskal-Wallis test followed by Dunn’s multiple comparisons test), indicating that superficial astrocytes are built from more compact loop elements rather than simply a greater number of larger structures.

Because LLSs have been linked to synaptic sites^37^, we next asked whether their superficial enrichment reflects a more synapse-facing organization. PSD-95 immunostaining showed that 71%–83% of LLSs across MOp layers either contained or directly apposed PSD-95 puncta, exceeding shuffled controls (61%; **Figures 2G, 2H, 2J**, **S2E** and **S2F**). Individual LLSs could also be associated with multiple PSD-95 puncta (**Figures 2I, 2K**, **S2C**, and **S2D**), indicating that a single loop element can interface with more than one excitatory postsynaptic structure. These data identify LLSs as synapse-associated modules of astrocyte fine processes and suggest that superficial astrocytes are especially enriched for these modules.

Finally, nearest-neighbor analysis showed that LLSs were positioned more closely together in L1 astrocytes than in deeper cortical layers or DLS (**Figures 2L** and **2M**; L1: 1.582 ± 0.063 µm, n = 12 ROIs, L2/3: 2.864 ± 0.708 µm, n = 12 ROIs, L5: 1.933 ± 0.082 µm, n = 12 ROIs, L6: 1.913 ± 0.087 µm, n = 12 ROIs; N = 3 mice; Kolmogorov-Smirnov test). Superficial astrocytes therefore combine a compact territorial footprint with high-density, small, and closely spaced synapse-associated loop elements. Together, these findings show that L1 astrocytes deploy a specialized superficial fine-process program rather than a scaled-down version of deeper-layer astrocytes, and they predict distinctive local signal integration, which we tested next by imaging spontaneous calcium activity.

### Superficial astrocytes exhibit disproportionately frequent spontaneous calcium activity

Because LLSs have been linked to local astrocytic calcium hotspots^37^, the dense and closely spaced LLS network of L1 astrocytes led us to ask whether superficial astrocytes also occupy a distinct functional state. We therefore imaged spontaneous calcium activity in sparsely labeled astrocytes from acute coronal slices of Aldh1l1-CreER^T2^;Lck-GCaMP6f mice^48^ across MOp layers and DLS using two-photon microscopy (**Figure 3A** and **S3B**). To capture the full spatiotemporal repertoire of events arising in both major branches and fine processes, we analyzed recordings with the event-based Astrocyte Quantitative Analysis (AQuA) framework^49–51^ rather than predefined static ROIs (**Figures 3B** and **3C**). Astrocytes in all regions displayed diverse spontaneous events during 5-min recordings (**Figure 3D**), but their activity profiles differed strikingly across depth.

**Figure 3.**
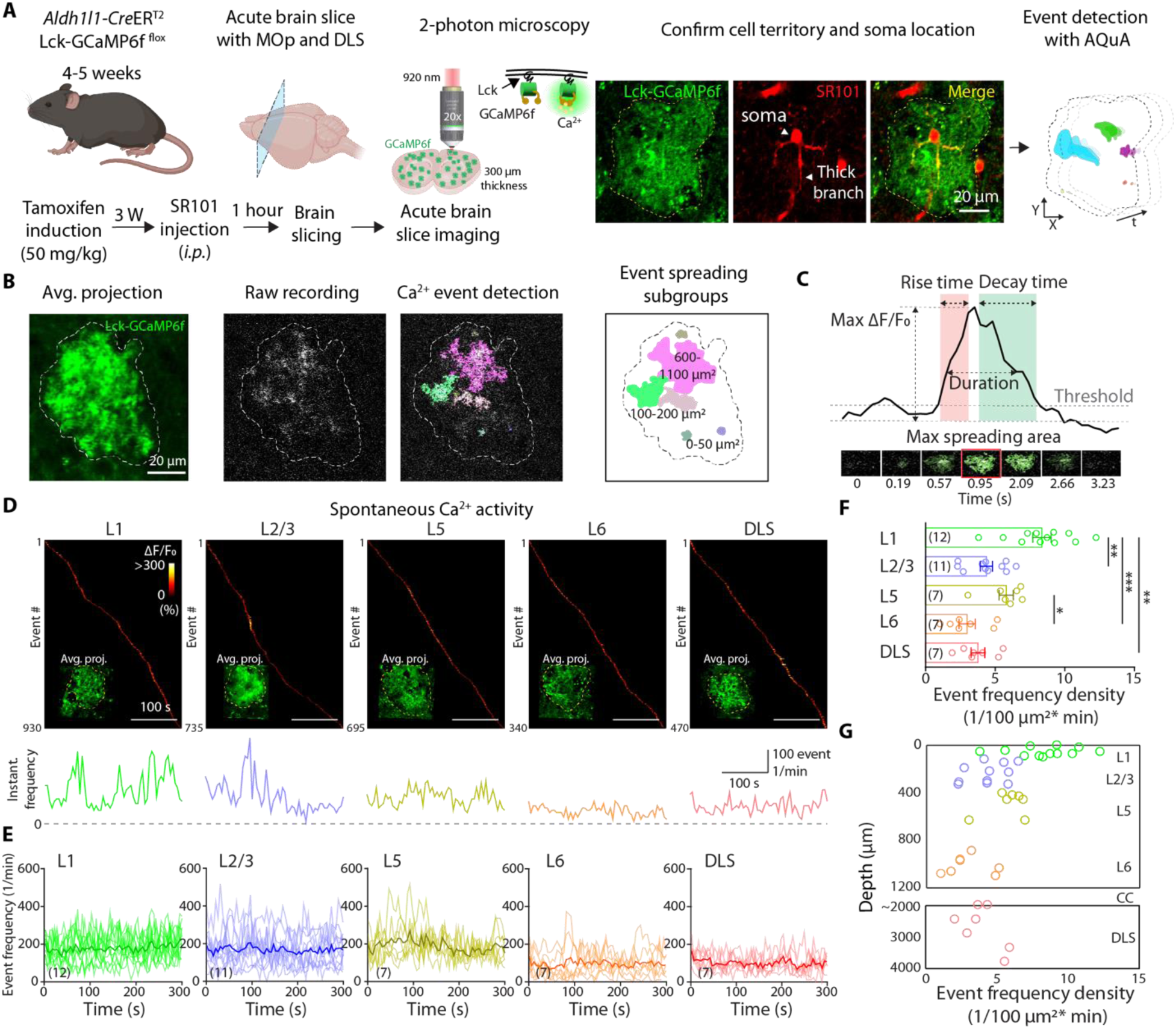
Heterogeneity in calcium activity across the MOp and DLS. (A) Experimental outline highlighting the steps from tamoxifen-induction to imaging analysis. (B) Astrocyte expressing Lck-GCaMP6f, exhibits diverse calcium events, detected using AQuA. The events are further subdivided into spreading subgroups. Scale bars represent 20 µm. (C) A typical calcium event trace showing the parameters used to make the comprehensive comparison across layers. (D) Heatmap showing the calcium events as they occur in a single astrocyte during the recording span of five minutes. Scale bars represent 100 seconds. Each heatmap has its event frequency plotted at the bottom. Colorbar represents the ΔF/F^0^ values range. (E) Event frequency plotted for each region; the thick line represents their mean trace. (F) Event frequency density across regions. (L1: 0.0835 ± 0.0066 events/10^2^ µm^2^ per minute, n = 12, L2/3: 0.0436 ± 0.0043 events/10^2^ µm^2^ per minute, n = 11, L5: 0.0579 ± 0.0050 events/10^2^ µm^2^ per minute, n = 7, L6: 0.0298 ± 0.0058 events/10^2^ µm^2^ per minute, n = 7, DLS: 0.0375 ± 0.0049 events/10^2^ µm^2^ per minute, n = 7; Kruskal-Wallis test followed by Dunn’s multiple comparisons test) (G) Plot showing the event frequency density across the astrocytes in the MOp and the DLS, along with their recording depth. The data are shown as mean ± SEM. *p < 0.05, **p < 0.01, ***p < 0.001.

Consistent with the structural enrichment of LLSs in superficial cortex, L1 astrocytes exhibited the highest event frequency density despite occupying the smallest territories (**Figures 3E–3G**). Event frequency density was greatest in L1 (8.4 ± 0.7 events/10^2^ µm^2^ per minute) and substantially lower in deeper cortex and DLS, reaching 3.0 ± 0.6 events/10^2^ µm^2^ per minute in L6 and 3.8 ± 0.5 events/10^2^ µm^2^ per minute in DLS (**Figure 3F**). This superficial bias was not limited to a single event class: when events were stratified by propagation size, L1 astrocytes remained the most active population across event-size bins (**Figure S3C**). Thus, the compact superficial astrocyte territory is not functionally sparse; rather, it supports the most active spontaneous signaling regime among the regions examined.

We next asked whether this elevated activity depends on ongoing action-potential firing. Acute tetrodotoxin (TTX, 1 µM) application did not significantly alter spontaneous event frequency in L1 and L2/3 astrocytes relative to baseline (**Figures S3D** and **S3E**), indicating that much of this activity persists after blockade of sodium-dependent spiking under these slice conditions, consistent with *in vivo* observation in rodent cortex^35^. Together, these results identify L1 astrocytes as a functionally distinct superficial population whose dense synapse-associated nanoarchitecture is matched by disproportionately frequent spontaneous calcium activity. This finding prompted us to next ask whether individual calcium events in superficial astrocytes are also specialized in their kinetics and spatial spread.

### Superficial astrocytes operate in a fast, spatially expansive calcium signaling mode

The elevated spontaneous activity of L1 astrocytes raised a key question: do superficial astrocytes simply generate more calcium events, or do they operate in a distinct signaling regime? To distinguish between these possibilities, we compared multiple AQuA-derived event features across MOp layers and DLS. Representative single-cell event maps from L1, L2/3, and L6 astrocytes already suggested qualitative differences in event organization beyond frequency alone (**Figures 4A–4C** and **S5**).

**Figure 4.**
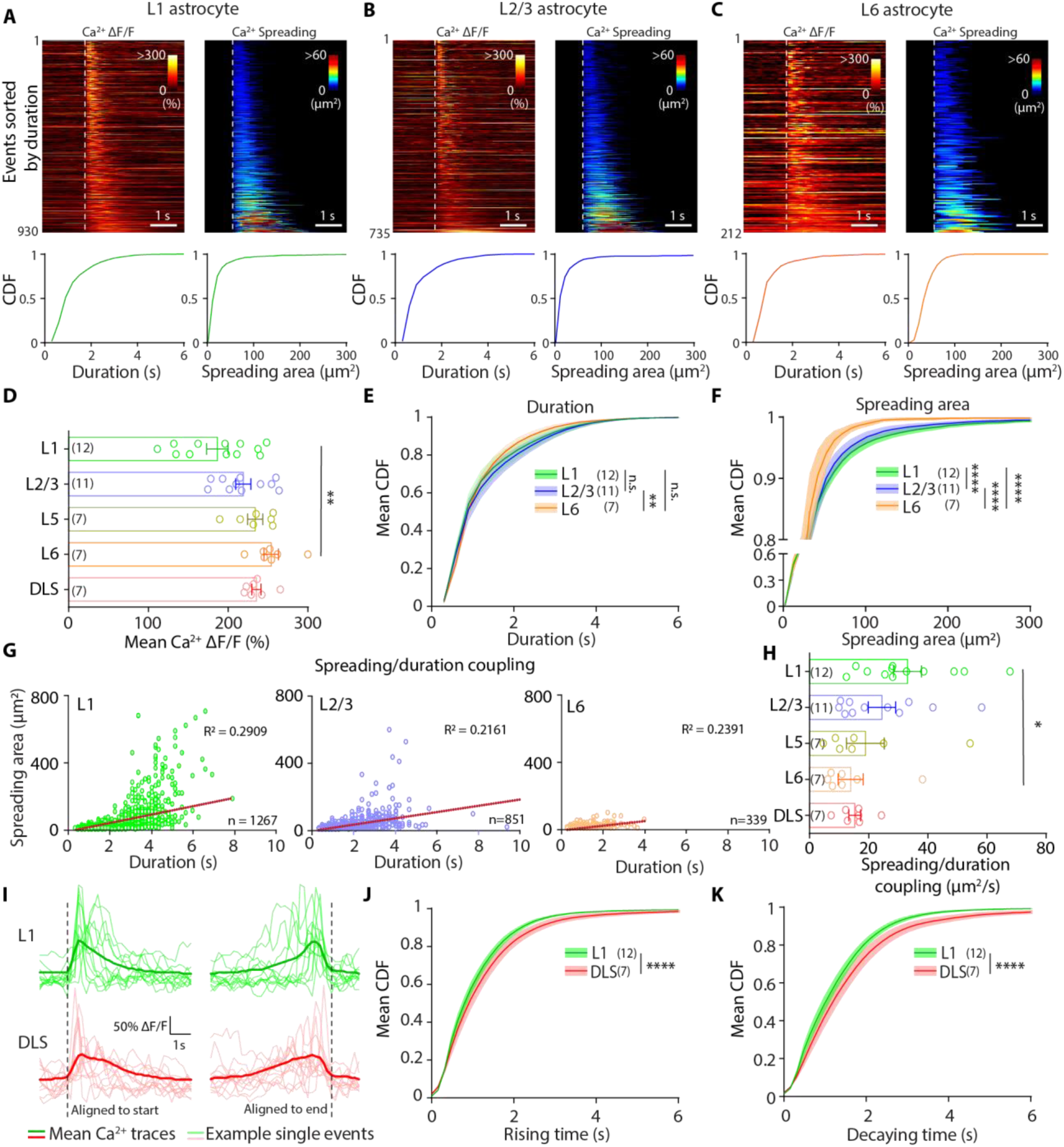
Calcium event parameter comparison of astrocytes across the MOp and DLS. (A) Calcium events occurring in a single astrocyte recorded from L1. Left: events sorted based in their duration, heatmap represents their amplitude range. Bottom: cumulative distribution of the duration of all events occurring in the astrocyte. Scale bar represents 1 second. Right: events sorted based in their duration, heatmap represents their spreading/propagation size (μm^2^). Bottom: cumulative distribution of the spreading size/propagation size of all events occurring in the astrocyte. Scale bar represents 1 second. (B) Calcium events occurring in a single astrocyte recorded from L2/3. Left: events sorted based in their duration, heatmap represents their amplitude range. Bottom: cumulative distribution of the duration of all events occurring in the astrocyte. Scale bar represents 1 second. Right: events sorted based in their duration, heatmap represents their spreading/propagation size (µm^2^). Bottom: cumulative distribution of the spreading size/propagation size of all events occurring in the astrocyte. Scale bar represents 1 second. (C) Calcium events occurring in a single astrocyte recorded from L6. Left: events sorted based in their duration, heatmap represents their amplitude range. Bottom: cumulative distribution of the duration of all events occurring in the astrocyte. Scale bar represents 1 second. Right: events sorted based in their duration, heatmap represents their spreading/propagation size (µm^2^). Bottom: cumulative distribution of the spreading size/propagation size of all events occurring in the astrocyte. Scale bar represents 1 second. (D) The mean Calcium ΔF/F_0_ of astrocytes recorded from the MOp and the DLS. (n = 12 (L1), n = 11 (L2/3), n = 7 (L5), n = 7 (L6), n = 7 (DLS) astrocytes; N = 19 mice; Kruskal-Wallis test followed by Dunn’s multiple comparisons test). (E) The comparison of the mean CDF of duration of L1, L2/3 and L6 astrocytes. (n = 12 (L1), n = 11 (L2/3), n = 7 (L6) astrocytes; N = 17 mice; Kolmogorov-Smirnov’s test). (F) The comparison of the mean CDF of spreading size of L1, L2/3 and L6 astrocytes. (n = 12 (L1), n = 11 (L2/3), n = 7 (L6) astrocytes; N = 17 mice; Kolmogorov-Smirnov’s test). (G) The spreading/duration coupling of calcium events in L1, L2/3 and L6 astrocytes. Each plot represents a single cell each. (Event number: 1267 (L1), 851 (L2/3), 339 (L6). (H) Quantification of the spreading/duration coupling of calcium events in the MOp and DLS astrocytes. (n = 12 (L1), n = 11 (L2/3), n = 7 (L5), n = 7 (L6), n = 7 (DLS) astrocytes; N = 19 mice; Kruskal-Wallis test followed by Dunn’s multiple comparisons test). (I) Representative traces showing the rising and decay time traces of L1 and DLS astrocytes, aligned to their starting and ending frame. 12 representative traces each are plotted from a L1 and a DLS astrocyte. The mean ± SEM (thick trace) is obtained from 929 (L1) and 434 (DLS) calcium events. (J) The comparison of the mean CDF of rising time of L1 and DLS astrocytes. (n = 12 (L1) astrocytes, N= 9 mice; n = 7 (DLS) astrocytes, N = 6 mice; Kolmogorov-Smirnov’s test). (K) The comparison of the mean CDF of decaying time of L1 and DLS astrocytes. (n = 12 (L1) astrocytes, N= 9 mice; n = 7 (DLS) astrocytes, N = 6 mice; Kolmogorov-Smirnov’s test). The data are shown as mean ± SEM. *p < 0.05, **p < 0.01, ****p < 0.0001.

Event amplitude did not explain the superficial phenotype. In fact, L1 astrocytes exhibited lower mean ΔF/F₀ than L6 astrocytes (**Figure 4D**), indicating that their elevated activity is not simply driven by larger fluorescence transients. Event duration alone also showed more limited layer selectivity, with L2/3 astrocytes displaying longer-lasting events than L6 astrocytes (**Figure 4E**). By contrast, spatial spread clearly distinguished the superficial population. L1 events propagated over larger areas than events in L2/3 or L6 astrocytes (**Figures 4F** and **S5**), and the largest L1 events approached the scale of nearly the entire astrocyte territory (**Figure S4A**). Because maximal event size in every region remained below the full cell area, this effect cannot be explained simply by differences in astrocyte territory size.

Instead, superficial astrocytes were distinguished by stronger coupling between event persistence and spatial recruitment. Plotting event duration against propagation revealed significantly higher spreading-duration coupling in L1 than in L6 astrocytes (33.1 ± 4.7 versus 13.9 ± 4.2 (µm^2^/ second); **Figures 4G** and **4H**), indicating that in superficial cells, events that last longer are also more likely to recruit a larger fraction of the astrocyte. This broad spatial engagement was accompanied by faster kinetics: compared with DLS astrocytes, L1 astrocytes showed steeper rise and decay phases and significantly shorter rise and decay times (**Figures 4I–4K** and **S6**).

Thus, the superficial phenotype is not simply an increase in event number. Rather, L1 astrocytes operate in a distinct spatiotemporal mode characterized by lower-amplitude, broader, faster, and more tightly coupled calcium events. Together with their dense synapse-associated LLS nanoarchitecture, these findings argue that superficial astrocytes are specialized to rapidly recruit large portions of a compact territory, a prediction we next tested by asking how calcium activity is distributed across the astrocyte as stable hotspot patterns.

### Heterogeneity in spontaneous calcium event hotspot pattern across astrocytes

Reports have shown that stable calcium hotspots are observed within individual astrocytes across different behavioral states and these are stable across days of training^21^. These stable calcium hotspots suggest that they are not random occurring events and might play a role in motor learning within the underlying neural network. The heterogeneity of the calcium events across regions made us question their hotspot patterns across these regions, whether there is any region-specific hotspot pattern exhibited by their spontaneous calcium events. We quantified the activated regions within each astrocyte territory. The event hotspot pattern across regions shows that L1, L2/3 and L5 astrocytes show hotspots that occupy most of their astrocytic territory (**Figure 5A**) while the L6 and DLS astrocytes tend to exhibit more localized/focal hotspots that are scattered within their territory (**Figure 5B-E**). Upon quantification we report that the L1 astrocytes have higher proportion of activated regions within their territory in comparison to the L6 and DLS astrocytes (**Figure 5F**; L1: 50.14 ± 6.821 %, n = 7, L2/3: 37.98 ± 6.725 %, n = 7, L5: 33.23 ± 8.269 %, n = 7, L6: 14.42 ± 4.138 %, n = 7, DLS: 19.11 ± 4.609 %, n = 7; Kruskal-Wallis test followed by Dunn’s multiple comparisons test). In order to see how the events occur within astrocytes, we also plotted their hotspot pattern across minutes of their recording. This quantification revealed that the L1 astrocytes exhibit higher territory activation across the entire recording (**Figure 5G** and **Figure S7**). It was interesting to note that the last minute of recording showed the highest event accumulation particularly in L1 astrocytes. It suggests that these cells are fast responsive to any kind of stress, in this case it might be photodamage, in comparison to other regions.

**Figure 5.**
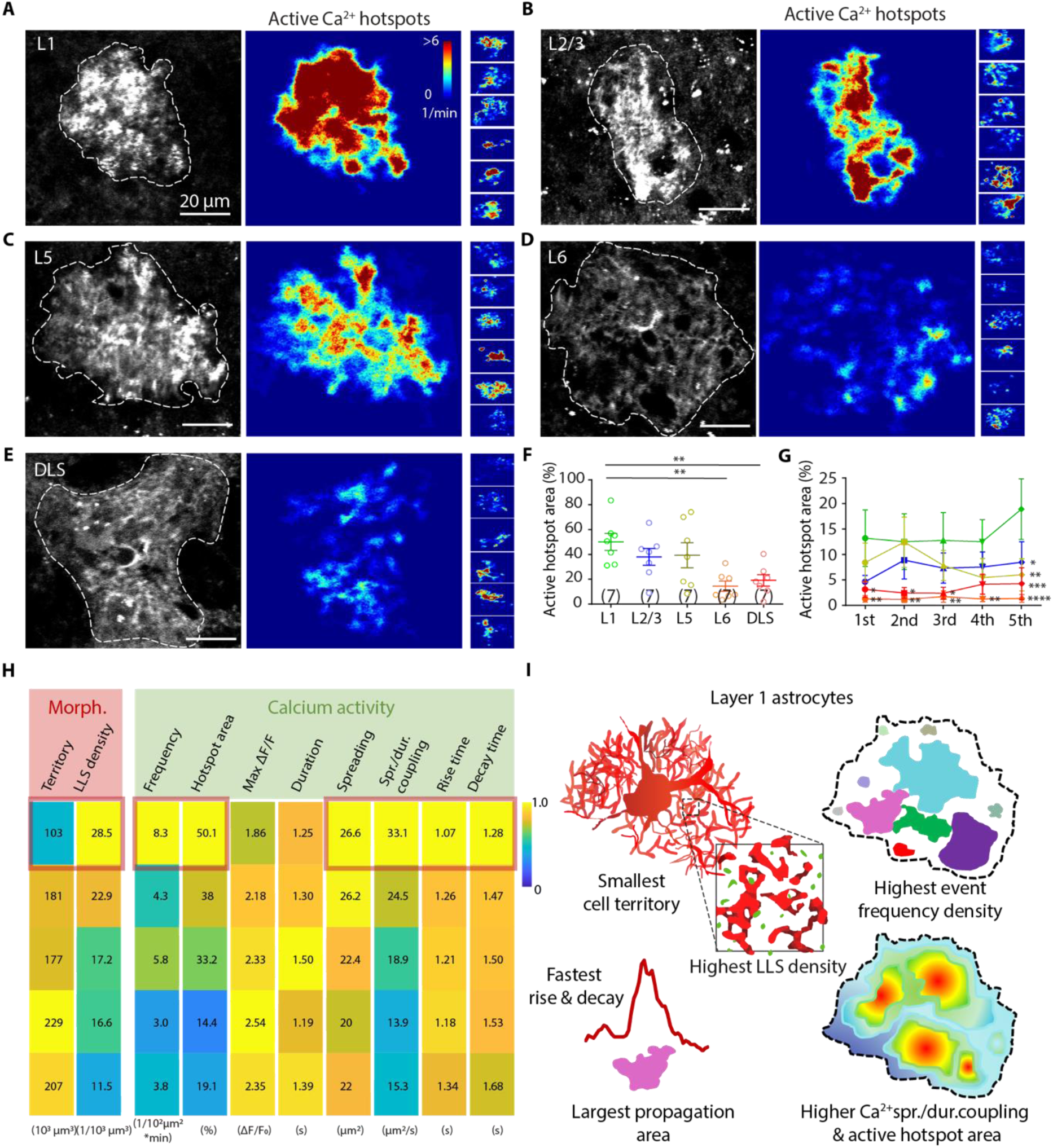
Event hotspot patterns show layer-wise differences. (A-E) Left: Average projection of L1 (A), L2/3 (B), L5 (C), L6 (D), and DLS (E) astrocytes. Astrocyte territory expressing Lck-GCaMP6f. Right: accumulated event patterns or hotspots of astrocytes obtained in 5 minutes of recording, from each brain regions. The heatmap represents the number of events occurring per minute within the astrocyte. The white arrows denote the event hotspots. (F) Quantification of the percentage of the activated territory region across the MOp and DLS astrocytes. (n = 7 (L1), n = 7 (L2/3), n = 7 (L5), n = 7 (L6), n = 7 (DLS) astrocytes; N = 14 mice; Kruskal-Wallis test followed by Dunn’s multiple comparisons test). (G) Quantification of the percentage of the activated territory region through every minute of recording, across the MOp and DLS astrocytes. (n = 7 (L1), n = 7 (L2/3), n = 7 (L5), n = 7 (L6), n = 7 (DLS) astrocytes; N = 14 mice; two-way ANOVA test followed by Tukey’s multiple comparisons test). The asterisks represent the comparisons of other regions with L1 astrocytes across minutes. (0-1 min; L1: 13.19 ± 5.537 %, n = 7, L2/3: 4.679 ± 1.240 %, n = 7, L5: 8.421 ± 3.827 %, n = 7, L6: 1.262 ± 0.649 %, n = 7, DLS: 3.143 ± 1.602 %, n = 7; 1-2 min; L1: 12.51 ± 5.462 %, n = 7, L2/3: 8.926 ± 3.700 %, n = 7, L5: 12.43 ± 4.946 %, n = 7, L6: 1.148 ± 0.589 %, n = 7, DLS: 2.427 ± 1.053 %, n = 7; 2-3 min; L1: 12.76 ± 5.466 %, n = 7, L2/3: 7.357 ± 2.974 %, n = 7, L5: 7.839 ± 2.971 %, n = 7, L6: 1.690 ± 1.124 %, n = 7, DLS: 2.375 ± 1.154 %, n = 7; 3-4 min; L1: 12.51 ± 4.329 %, n = 7, L2/3: 7.581 ± 2.958 %, n = 7, L5: 5.442 ± 3.749 %, n = 7, L6: 1.235 ± 0.494 %, n = 7, DLS: 4.140 ± 1.961 %, n = 7; 4-5 min; L1: 18.90 ± 5.934 %, n = 7, L2/3: 8.506 ± 4.028 %, n = 7, L5: 6.013 ± 3.140 %, n = 7, L6: 1.347 ± 0.770 %, n = 7, DLS: 4.241 ± 3.133 %, n = 7, two-way ANOVA test followed by Tukey’s multiple comparisons test). (H) Table summarizing the comparison of all the event parameters studied across the different layers of the MOp and DLS. (I) Illustration highlighting the distinct morphological and calcium activity properties of the L1 astrocytes. The data are shown as mean ± SEM. *p < 0.05, **p < 0.01, ***p < 0.001, ****p < 0.0001.

In summary, we found that the L1 astrocytes are the most active subpopulation within the MOp. They have the smallest territory size, high LLS density, highest event frequency density and a high activation region. In addition, they also exhibit large spreading events, showing coupling with the event duration. These astrocytes exhibit shorter event duration, fastest rising and decay time but large spreading area, this indicates that might indeed play a role in faster integration of inputs across synapses (**Figure 5H**). The illustration summarizes the morphological and calcium activity properties’ heterogeneity observed across the MOp and the DLS, highlighting the distinct properties of the L1 astrocytes (**Figure 5I**).

### Identification and study of L1 astrocyte gene expression profile from publicly available scRNA-seq dataset

The heterogeneity in morphology and calcium activity of MOp astrocytes led us to dig into the putative players that might be responsible for maintaining the high calcium activity of L1 astrocytes. In order to understand the underlying mechanism, we looked into the publicly available scRNA-seq dataset of the MOp generated by the Brain Initiative Cell Census Network (BICCN)^1,52^ (**Figure 6A**). The UMAP plot shows the presence of distinct clusters occupied by different subtypes of neuronal and non-neuronal cells. The cluster enriched in astrocytes was chosen based on minimal expression of *Dcx* (**Figure 6B**) and high expression of pan astrocytic marker *Sox9* (**Figure 6C**). This cluster was thereby analyzed to generate the UMAP plot showing the presence of subpopulations of astrocytes within the MOp (**Figure 6D**). The dataset used in this study is specifically enriched in astrocytes as can be seen from the uniform expression of pan astrocytic markers *Aldh1l1* and *Sox9* (**Figure S8A** and **S8B**) across all the clusters.

**Figure 6.**
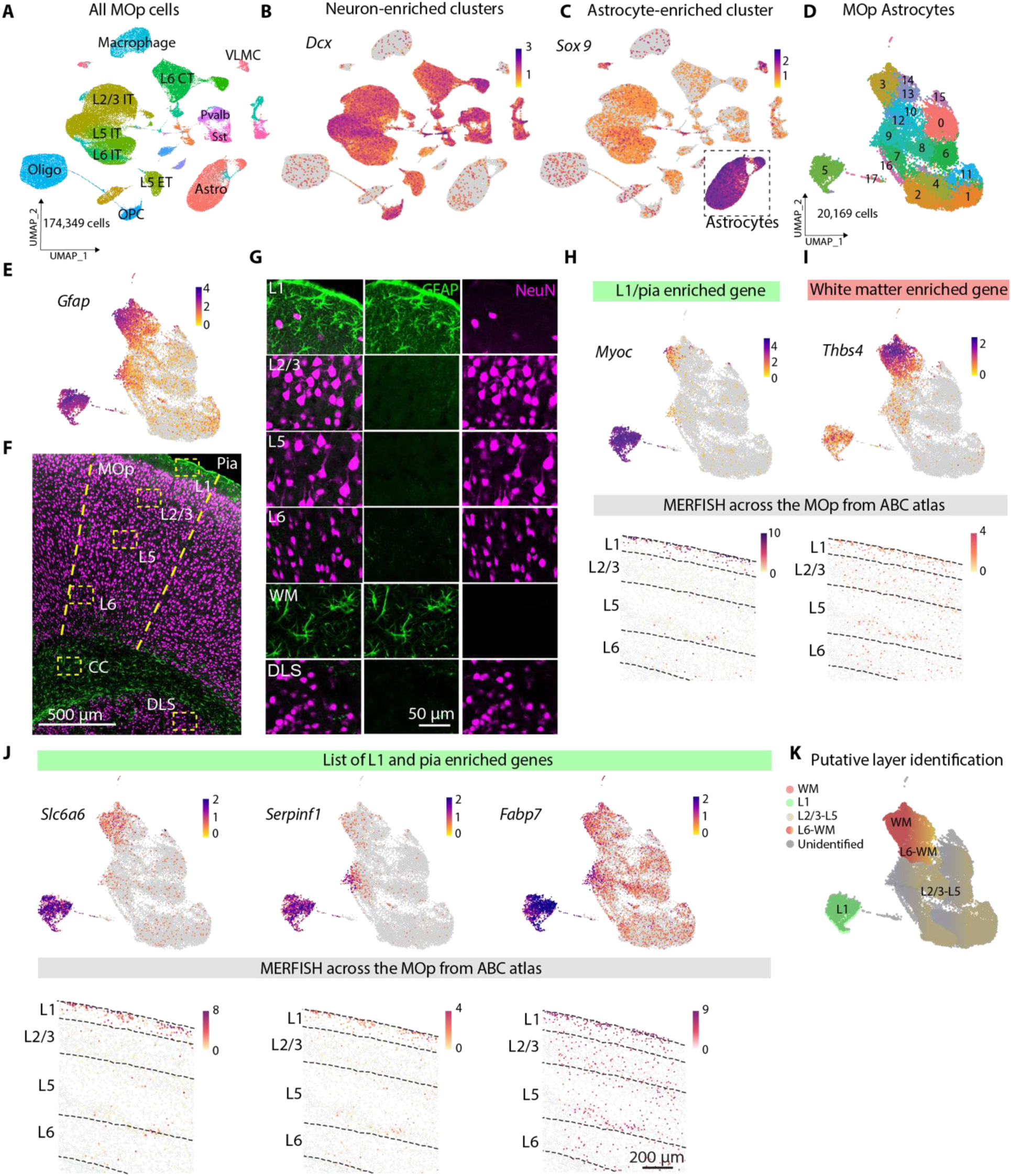
Astrocyte subpopulations in MOp revealed by scRNA-seq. (A) Single-cell RNA sequencing data of the primary motor cortex (MOp) from adult mouse brain generated by the Brain Initiative Cell Census Network (BICCN)^1^. It shows the different clusters occupied by neuronal and non-neuronal cells. (174,349 cells). Oligo: oligodendrocyte, VLMC: vascular leptomeningeal cell, Astro: astrocyte, OPC: oligodendrocyte progenitor cell, ET: extratelencephalic, IT: intratelencephalic, CT: corticothalamic, Pvalb: parvalbumin, Sst: somatostatin. (B) UMAP plot showing the enrichment of *Dcx* in the neuronal clusters. (C) UMAP plot showing the enrichment of *Sox9* in the astrocyte cluster. (D) UMAP generated from analysis of the astrocyte cluster showing 18 sub clusters. (20,169 cells) (E) UMAP showing *Gfap* enrichment in cluster 3 and 5. (F) Overview of GFAP (green) expression in the MOp astrocytes. The yellow rectangles correspond to regions within the different layers and the dorsolateral striatum (DLS). Scale bars represent 500 μm. (G) Layer-wise and DLS expression of GFAP (green) and NeuN (magenta) within the yellow rectangles. Scale bars represent 50 μm. (H) UMAP showing enrichment of *Myoc* (L1/pia enriched marker) in cluster 5 (top). MERFISH data from the ABC atlas (https://doi.org/10.35077/g.610) showing the spatial distribution of *Myoc* (bottom). (I) UMAP showing enrichment of *Thbs4* (WM enriched marker) in cluster 3 (top). MERFISH data from the ABC atlas (https://doi.org/10.35077/g.610) showing the spatial distribution of *Thbs4* (bottom). (J) List of genes enriched in the L1 and pia. (K) Putative layer-specific identification of astrocytic clusters in the MOp.

In order to identify the L1 astrocyte cluster, we first looked into the well-known astrocytic marker *Gfap*. This gene was highly enriched in clusters 3 and 5 (**Figure 6E**). Identification of these clusters were done by performing immunohistochemistry (**Figure 6F**). The L1 astrocytes showed a clear high expression of *Gfap*. This level starts to decrease gradually as we move down into the L2/3 and the L5. However, the expression starts to increase again from deeper L6 to the white matter (WM). The expression of GFAP is observed solely in astrocytes, as can be observed from the absence of any overlap between GFAP and NeuN in neurons (**Figure 6G**). We then came across the genes *Myoc* and *Thbs4*^53^ which are reported to be a pial astrocyte marker and a gene specific to white matter astrocytes, respectively. The UMAP plots clearly shows confined enrichment of the cluster 5 with *Myoc* and cluster 3 with *Thbs4* (**Figures 6H** and **6I**), suggesting that they are L1 and WM astrocytes enriched cluster, respectively. Further confirmation of the L1 cluster was done by additional genes (**Figure S8C**), that showed clear high expression in the MERFISH^52^ data (<u>Brain Knowledge Platform</u>) of the L1 astrocytes in the MOp (**Figures 6J and S8C**). This also confirmed that the cluster 5 is a mixture of both pial and L1 astrocytes.

Based on the survey of genes^3,11,54^ available in previous reports, we found that these clusters are very heterogeneous in their expression patterns of different genes. Unlike report in the SSp^3^, the MOp astrocytes do not show clear expression enrichment of genes in the L2/3 and L5 astrocytes, therefore, identification of the other clusters is currently incomplete (**Figure S9**). We provide a putative layer identification of the astrocyte clusters, although further validation is highly essential (**Figure 6K, S8C**). Notably, the genes enriched in cluster 5 included candidates associated with extracellular matrix modification (*Sulf2*^55,56^, *Myoc*^57,58^), guidance signaling (*Nrp2*, *Sema5a*)^59,60^, and metabolic transport (*Fabp7*^61–63^, *Slc38a1*^64,65^), suggesting that L1/pia astrocytes are supported by a distinct structural and metabolic program (**Figure 6K, S8C**).

### Layer 1 astrocyte morphology and calcium activity likely regulated by *Id1* and *Id3* transcriptional regulators

GFAP is a widely used astrocyte marker, however, this protein shows varying expression levels across different regions. It has been reported that in Alexander’s disease, astrocytes show high GFAP levels along with ‘extra-large’ calcium signals, which made us question if there is a correlation between them^66^. To test this hypothesis, we also recorded astrocytic spontaneous calcium activity from the hippocampus CA1 area and primary visual cortex-Layer1 (VISp-L1) which show high and low expression of GFAP^54^, respectively (**Figures S10A** and **S10B**). Our observations indeed support this correlation, CA1 astrocytes tend to show high event frequency density while the VISp-L1 astrocytes exhibit a very low event frequency density (**Figures S10C**). There is indeed a correlation between astrocytic GFAP levels and their calcium event frequency density (**Figure S10D**).

Along with *Gfap*, we found two other genes *Id1* and *Id3* that show similar expression pattern (**Figure 7A)**. It has been reported that Id1, Id3 plays a key role in promoting astroglial differentiation by inhibiting neuronal differentiation and thereby facilitating expression of genes like *Gfap* and *S100β*^46^. These proteins are highly essential in this switch to astroglia^67–69^. There is direct evidence of *Gfap* being regulated by Id proteins and indirect correlation between astrocytic GFAP levels and their calcium event frequency density. However, if these Id proteins also regulate astrocytic calcium activity, is not known. Therefore, we set out to ask if the sustained *Id1*, *Id3* gene levels in adult L1 astrocytes in the MOp plays any role in maintaining their distinct morphology and heightened calcium activity. The expression of Id3 in L1 astrocytes was confirmed by using histology studies, with Sox9 as a comparison which is expressed by most astrocytes across the cortex (**Figure 7B)**. The intensity plot shows the high expression of Id3 in the L1 astrocytes while the Sox9 shows a more similar intensity across layers (**Figure 7C**; L1: 139.8 ± 12.2 a.u., n = 8, L2/3: 79.91 ± 10.75 a.u., n = 8, L5: 73.24 ± 9.937 a.u., n = 8, L6: 75.77 ± 9.416 a.u., n = 8, and **Figure 7D**; L1: 249.0 ± 0.8804 a.u., n = 8, L2/3: 245.5 ± 1.018 a.u., n = 8, L5: 244.4 ± 1.701 a.u., n = 8, L6: 246.3 ± 1.548 a.u., n = 8, Kruskal-Wallis test followed by Dunn’s multiple comparisons test).

**Figure 7.**
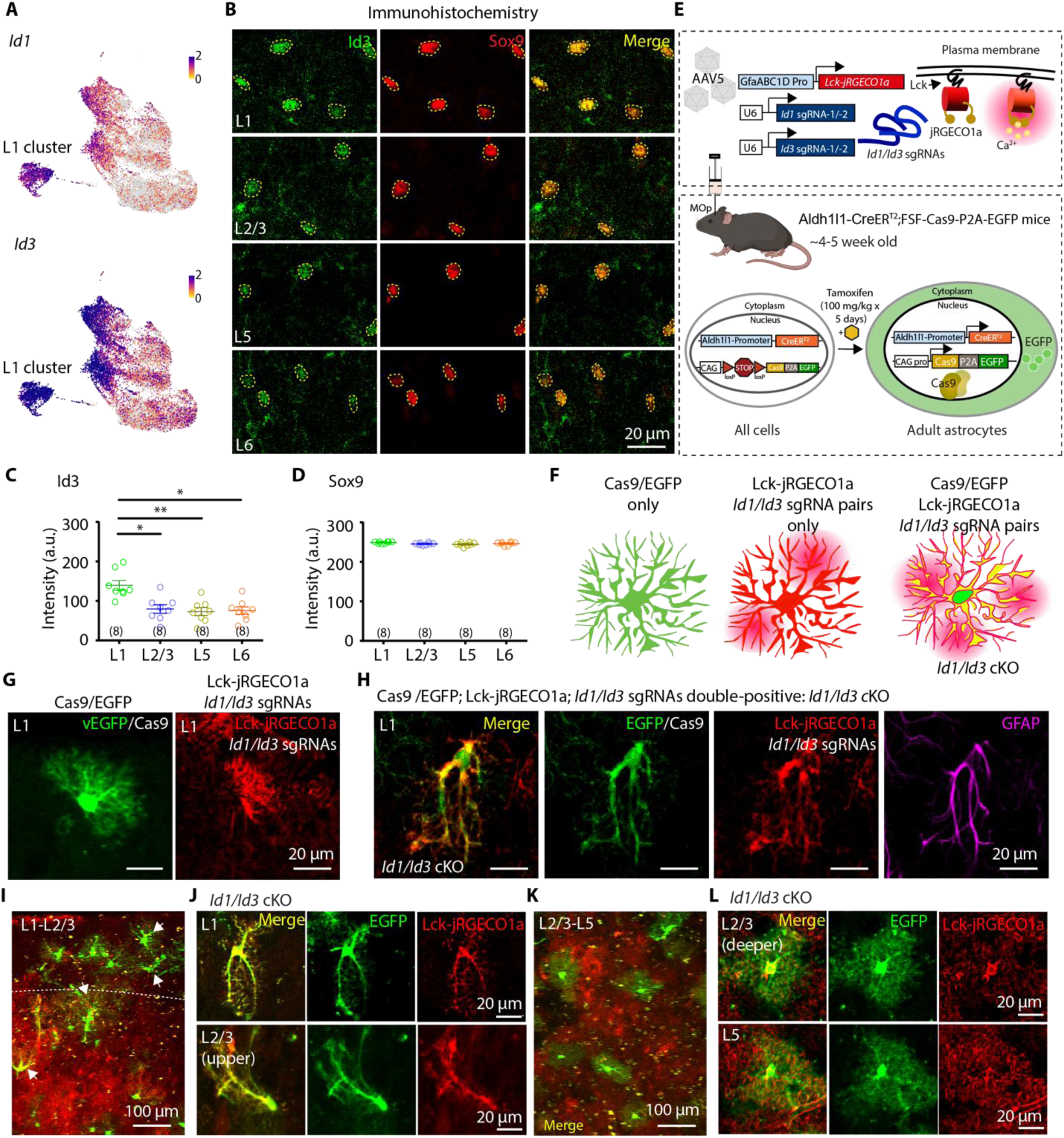
Conditional knockout of *Id1* and *Id3* genes lead to abnormal morphology in L1 astrocytes. (A) UMAPs showing the expression pattern of *Id1* and *Id3* genes. (B) Confocal images of Id3 (green), Sox9 (red) and their co-staining (yellow) pattern across layers in the MOp. Scale bars represent 20 µm. The yellow dashed lines mark the outline of the astrocyte somata. (C) The intensity of Id3 across the astrocytes in the MOp. (n = 8 (L1), n = 8 (L2/3), n = 8 (L5), n = 8 (L6) slices; N = 4 mice; Kruskal-Wallis test followed by Dunn’s multiple comparisons test). (D) The intensity of Sox9 across the astrocytes in the MOp. (n = 8 (L1), n = 8 (L2/3), n = 8 (L5), n = 8 (L6) slices; N = 4 mice; Kruskal-Wallis test followed by Dunn’s multiple comparisons test). (E) Experimental outline highlighting the steps for conditional knockout (cKO) strategy of *Id1* and *Id3* genes using CRISPR-Cas9 technology. (F) Illustration showing all the possible morphological outcomes. (G) Astrocytes expressing only Cas9-EGFP (left) or only Lck-jRGECO1a (right). (H) Astrocytes after cKO, expressing Cas9-EGFP (green), Lck-jRGECO1a (red), GFAP (magenta). Scale bars represent 20 µm. (I) Overview of the L1-L2/3 region in the MOp of mice expressing Cas9-EGFP and Lck-jRGECO1a in astrocytes. Scale bars represent 100 µm. (J) cKO phenotype of L1 astrocytes expressing both Cas9-EGFP (green) and Lck-jRGECO1a (red) (top). cKO phenotype of upper L2/3 astrocytes expressing both Cas9-EGFP (green) and Lck-jRGECO1a (red) (bottom). Scale bars represent 20 µm. (K) Overview of the L2/3-L5 region in the MOp of mice expressing Cas9-EGFP and Lck-jRGECO1a in astrocytes. Scale bars represent 100 µm. (L) cKO phenotype of deeper L2/3 astrocytes expressing both Cas9-EGFP (green) and Lck-jRGECO1a (red) (top). cKO phenotype of L5 astrocytes expressing both Cas9-EGFP (green) and Lck-jRGECO1a (red) (bottom). Scale bars represent 20 μm. The data are shown as mean ± SEM. *p < 0.05, **p < 0.01.

### Effect of *Id1*, *Id3* conditional knockout on adult MOp astrocytic morphology

To explore the role of these transcriptional regulators in adult MOp astrocytes, we designed a CRISPR-Cas9 mediated knockout strategy^70^. We designed plasmid constructs to express single-strand guide RNA pairs against genes *Id1* and *Id3* and Lck-jRGECO1a, a membrane-tethered red calcium indicator. The Lck-jRGECO1a is used as both an indicator of the expression of the guide RNAs and the recording of spontaneous calcium activity from astrocytes. This construct was packaged into AAV5 and injected into the MOp of Aldh1l1-CreER^T2^;LSL-Cas9-EGFP mice (**Figure 7E)**. These mice are further induced by tamoxifen (100 mg/kg-5 consecutive days) to obtain sparse labeling of astrocytes expressing Cas9-EGFP. The illustration shows all the combinations of the expected phenotypes, astrocytes might express either only Cas9-EGFP or Lck-jRGECO1a, or both of these genes (**Figure 7F).** The astrocytes expressing both Lck-jRGECO1a and EGFP are further recorded for the conditional knockout (cKO) study.

The astrocytes expressing only Cas9-EGFP or Lck-jRGECO1a, displayed intact morphology (**Figure 7G).** On the other hand, L1 astrocytes were highly affected by this cKO, showing an aberrant morphology but still expressing GFAP (**Figure 7H**). This suggests that *Gfap* expression in adult astrocytes might not be under direct regulation by Id proteins. These astrocytes display thicker primary branches but their fine processes are reduced (**Figure 7I** and **7J, Figures S11A** and **S11B**). They do not display the typical ‘cloudy’ or ‘spongiform’ morphology of astrocytes (**Figure S11D)**. The upper L2/3 astrocytes were also found to display the similar aberrant morphology (**Figure 7I** and **7J, Figures S11C)**. However, as we go deeper into the L2/3 and L5, the cKO effects on astrocyte morphology is not observed (**Figure 7K** and **7L, Figure S11D**). These astrocytes still display normal morphology despite cKO.

### Effect of *Id1*, *Id3* conditional knockout on adult MOp astrocytic spontaneous calcium activity

Simultaneous recording of the spontaneous intracellular calcium activity of the only Lck-jRGECO1a expressing astrocytes and the cKO ones expressing both Lck-jRGECO1a and Cas9-EGFP was performed (**Figures 8A** and **8B**). The L1 astrocytes expressing only Lck-jRGECO1a exhibited normal calcium event frequency density while the cKO astrocyte showed drastic reduction (**Figure 8C** and **8E**; L1: 6.905 ± 0.4956 events/10^2^ µm^2^ per minute, n = 5, L1-cKO: 1.845 ± 0.4558 events/10^2^ µm^2^ per minute, n = 9, Mann-Whitney U test). On the other hand, the L2/3 astrocytes show divided response upon cKO, with the deeper L2/3 astrocytes showing no effect (**Figure 8D** and **8F**; L2/3: 3.512 ± 0.6567 events/10^2^ µm^2^ per minute, n = 5, L2/3-cKO: 1.996 ± 0.6164 events/10^2^ µm^2^ per minute, n = 6, Mann-Whitney U test). Upon plotting all the cells recorded across the L1 and L2/3 of the MOp, we can see a shift to the left in event frequency density upon cKO in L1 astrocytes (**Figure 8G**). These results suggest that cKO of *Id1* and *Id3* genes may have an indirect effect on the L1 astrocyte morphology and calcium activity. The L1 astrocytes are positioned with high proximity to the pial surface, hence, they might require a stronger regulation to maintain their structure and function in the face of various adversities like mechanical, oxidative and inflammatory stresses at the blood-brain barrier (BBB). Because of these stresses the L1 astrocytes might be more vulnerable to fate changes. The heightened expression levels of these genes in the adult L1 astrocytes might play a role in maintaining their structure and function.

**Figure 8.**
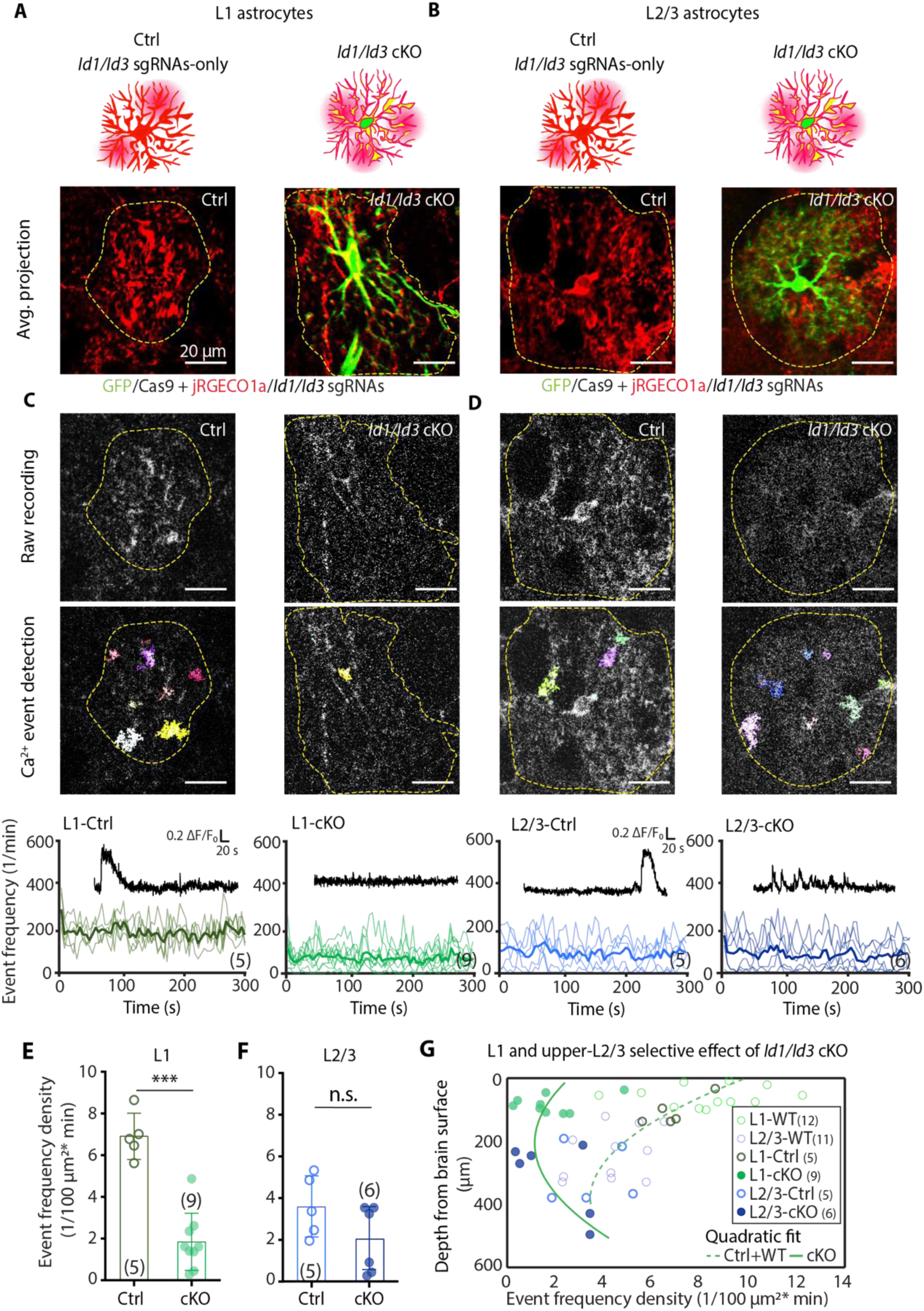
Knockout of *Id1* and *Id3* genes lead to aberrant spontaneous calcium activity in L1 astrocytes. (A) Top: Illustration showing the morphology of Ctrl and *Id1*/*Id3* cKO in L1 astrocytes. Bottom: Average projection of L1 astrocyte upon Ctrl and *Id1*/*Id3* cKO conditions. (B) Top: Illustration showing the morphology of Ctrl and cKO in L2/3 astrocytes. Bottom: Average projection of L2/3 astrocyte upon Ctrl and *Id1*/*Id3* cKO conditions. (C) Top, left: Raw recording of Ctrl L1 astrocyte expressing Lck-jRGECO1a. Top, right: Raw recording of cKO L1 astrocytes expressing Lck-jRGECO1a. Middle: event detection using AQuA in Ctrl and *Id1*/*Id3* cKO astrocytes. Bottom, left: event frequency plotted for L1-Ctrl; the thick line represents their mean trace. Inset: example of Ca^2+^ trace in L1-Ctrl condition. Bottom, right: event frequency plotted for L1-cKO; the thick line represents their mean trace. Inset: example of Ca^2+^ trace in L1-cKO condition. (D) Top, left: Raw recording of Ctrl L2/3 astrocyte expressing Lck-jRGECO1a. Top, right: Raw recording of *Id1*/*Id3* cKO L2/3 astrocytes expressing Lck-jRGECO1a. Middle: event detection using AQuA in Ctrl and *Id1*/*Id3* cKO astrocytes. Bottom, left: event frequency plotted for L2/3-Ctrl; the thick line represents their mean trace. Inset: example of Ca^2+^ trace in L2/3-Ctrl condition. Bottom, right: event frequency plotted for L2/3-cKO; the thick line represents their mean trace. Inset: example of Ca^2+^ trace in L2/3-cKO condition. (E) Event frequency density comparison between Ctrl and *Id1*/*Id3* cKO L1 astrocytes. (n = 5 (L1), n = 9 (cKO); N = 4 mice; two-tailed, Mann-Whitney U test). (F) Event frequency density comparison between Ctrl and *Id1*/*Id3* cKO L2/3 astrocytes. (n = 5 (L2/3), n = 6 (cKO); N = 4 mice; two-tailed, Mann-Whitney U test). (G) Summary plot of event frequency density vs the depth of Ctrl and *Id1*/*Id3* cKO astrocytes recorded from the MOp. The solid and dashed lines represent a quadratic fit used to visualize the trend in the control vs cKO Calcium recordings. (n = 12 astrocytes (L1-WT), N = 9 mice; n = 11 astrocytes (L2/3-WT), N = 9 mice; n = 5 astrocytes (L1-Ctrl), N = 2 mice; n = 9 astrocytes (L1-cKO), N = 4 mice; n = 5 astrocytes (L2/3-Ctrl), N = 2 mice; n = 6 astrocytes (L2/3-cKO), N = 2 mice). The data are shown as mean ± SD. ***p < 0.001, ns = non-significant.

## Discussion

Astrocytes, once regarded as a relatively homogeneous glial population, are now recognized as highly heterogeneous, exhibiting region- and layer-specific diversity in morphology, gene expression profiles, and physiology. In this study, we aimed to provide a comprehensive characterization of astrocytic heterogeneity in the MOp. By integrating super-resolution structural imaging, spontaneous calcium imaging, and analysis of publicly available scRNA-seq datasets, we demonstrate that astrocytes in the MOp differ markedly across cortical layers in their territory occupancy, fine-process architecture, calcium activity properties, and gene expression profiles.

We first observed pronounced layer-dependent differences in astrocytic morphology. Among all layers, L1 astrocytes occupied the smallest territorial volumes. While astrocytic morphological heterogeneity has previously been reported in the primary somatosensory cortex^4^, we found that in the MOp, astrocytes in L2/3 and L6 did not differ significantly in territory size or orientation angle. These findings suggest that astrocytic morphology is strongly influenced by the local extracellular and circuit environment. L1 lacks neuronal somata and is densely populated by horizontal apical dendritic tufts, receiving inputs from deeper cortical layers and neighboring brain regions. This unique organization likely constraints astrocytic territory size while promoting intimate interactions with dendritic spines. In contrast, deeper layers contain densely packed neuronal somas, thick apical and basal dendrites, and axonal bundles in the DLS, contributing to the larger territorial coverage of astrocytes in these regions. Consistent with previous reports, astrocytes in L2/3 and L5 can contact up to 8-10 neuronal somata, while striatal astrocytes adopt specialized morphologies to ensheath axonal bundles. Although astrocytic orientation angles are known indicators of injury or pathology, their functional relevance in the healthy brain such as whether they facilitate integration across layers or multiple neurons, remains unclear.

Astrocytic structural complexity represents another key dimension of heterogeneity. Astrocytes possess highly branched arbors, with fine processes constituting approximately 75% of their structure^47^. Using SR-SIM imaging, we visualized LLSs within astrocytic fine processes and quantified their distribution across layers. Previously, LLSs have been reported near synapses and associated with calcium hotspots^37^. Our data reveal a strong layer dependence in LLS density, with L1 astrocytes exhibiting the highest density, which progressively decreases in deeper layers such as L6 and DLS astrocytes. Co-localization analysis with PSD-95 confirmed that these structures are positioned in close proximity to synapses. The spatial closeness of LLSs in L1 astrocytes might play a key role in faster spatial integration of calcium events that occur in these LLSs. This also highlights an important question, whether this morphological specialty gives any advantage to L1 astrocytes’ calcium activity responses. The combination of small territorial volume and high LLS density in L1 astrocytes suggests an enhanced capacity to engage with densely packed synaptic inputs. This interpretation is consistent with previous reports showing layer-dependent differences in dendritic spine dynamics and synaptic ensheathment^4,71^, although such data are not yet available for the MOp.

Given that LLS and astrocytic fine processes are known sites of calcium signaling^72–74^, we next examined whether structural heterogeneity is reflected in spontaneous calcium activity. L1 astrocytes exhibited the highest calcium event frequency density, which declined gradually with cortical depth. Analysis of multiple event parameters revealed distinct layer-specific calcium signatures. L1 astrocytes displayed larger activation zones that occupied nearly half of their territory, faster rise and decay kinetics, broader spatial spreading, and longer event durations. These properties suggest that L1 astrocytes operate on faster timescales and may integrate inputs across multiple synapses simultaneously. Interestingly, L1 astrocytes showed lower calcium event amplitudes (ΔF/F₀), which may reflect elevated baseline calcium levels due to sustained activity. In contrast, deeper-layer astrocytes, particularly in L6, exhibited more localized, higher-amplitude events, consistent with their lower baseline activity. Across all layers, focal calcium events constituted the largest fraction of detected events, in agreement with prior studies. Differences in calcium dynamics may also be shaped by intracellular organelle composition, such as variations in mitochondrial or endoplasmic reticulum abundance across layers, although this remains to be tested.

Astrocytic heterogeneity is further reinforced at the transcriptional level. Multiple studies have reported region- and layer-specific astrocytic gene expression profiles^2,6,11^. Analysis of a publicly available MOp scRNA-seq dataset^1^ revealed that L1 astrocytes are enriched for *Gfap*, *Id1*, and *Id3*, among other genes. *Id1* and *Id3* are inhibitors of bHLH transcription factors and play key roles in promoting astroglial identity during development by suppressing neuronal differentiation and enabling expression of astrocytic markers such as *Gfap* and *S100β*^46,67–69^. Although these genes are well studied in developmental contexts, their role in maintaining astrocyte identity and function in the adult brain remains poorly understood.

Beyond *Gfap*, *Id1*, and *Id3*, the transcriptomic profile of cluster 5, the L1/pia enriched astrocyte cluster suggests a coordinated specialization program. Genes enriched in this cluster include extracellular matrix-related factors (*Sulf2*^55,56^, *Myoc*^57,58^), guidance-associated molecules (*Nrp2, Sema5a*)^59,60^, and transporters such as *Fabp7*^61–63^, *Slc38a1*^64,65^, and *Slc6a6*^75^. Together, these genes suggest that L1 astrocytes are specialized not only for structural interactions at the cortical surface but also for the metabolic support and niche responsiveness required for intense synaptic monitoring. Many of these genes also contain predicted E-box motifs, raising the possibility that they are regulated, directly or indirectly, by the Id-E-protein axis.

To address this, we performed CRISPR–Cas9–mediated knockout of *Id1* and *Id3* in adult astrocytes. Loss of these genes led to a striking phenotype in L1 astrocytes, characterized by expanded territorial volume, reduced fine-process complexity, and markedly diminished calcium activity. Similar effects were observed in superficial L2/3 astrocytes but not in deeper L2/3, L5, or L6 astrocytes, which retained normal morphology and function. These findings indicate a layer-specific dependence on *Id1* and *Id3*. Notably, GFAP expression was not abolished following knockout, suggesting that Id proteins may play distinct regulatory roles in adult astrocytes beyond lineage specification. Given the close association of L1 astrocytes with the pial surface, vasculature, and cerebrospinal fluid, these cells are exposed to diverse extrinsic signals that could destabilize their identity in the absence of intrinsic transcriptional regulators. *Id1* and *Id3* may therefore act to stabilize L1 astrocyte identity under such conditions. Future studies using scRNA-seq and ATAC-seq will be necessary to elucidate the downstream targets and epigenetic mechanisms underlying this regulation.

This framework suggests a structural-metabolic synergy downstream of *Id1* and *Id3*. Genes such as *Fabp7*^61–63^, *Myoc*^57,58^, and *Emb*^76^ may provide the lipid-handling, adhesive, and membrane-organizing capacity needed to build and maintain the dense loop-like structures of L1 astrocytes, whereas *Sulf2*^55,56^, *Nrp2*, and *Sema5a*^59,60^ may help these cells sense and remodel the extracellular niche at the cortical surface. In parallel, transporters such as *Slc38a1*^64,65^ and *Slc6a6*^75^ may support the metabolic and osmotic demands imposed by frequent calcium activity. We therefore propose that *Id1* and *Id3* may help sustain a specialized surface-astrocyte program by relieving E-protein-mediated repression of genes required for LLS formation, hyperactive calcium signaling, and synaptic surveillance. One intriguing possibility is that *Sulf2*-mediated extracellular matrix remodeling could further reinforce this superficial identity by modulating the local availability of instructive cues, potentially including BMP signals, near the pial surface.

In summary, this study provides a comprehensive, multi-modal characterization of astrocyte heterogeneity in the MOp. L1 astrocytes emerge as a uniquely specialized population distinguished by compact morphology, dense fine-process architecture, heightened and fast calcium signaling. Their selective dependence on *Id1* and *Id3* highlights the importance of layer-specific transcriptional programs in shaping astrocyte structure and function. Together, these findings support a model in which astrocytes play active, layer-specific roles in modulating neuron-astrocyte interactions and supporting local circuit dynamics, rather than serving as passive support cells.

## Acknowledgements

The authors thank Dr. Yijuang Chern, Dr. Cheng-Ting Chien, Dr. Sheng-Hong Chen, Dr. Suewei Lin, Dr. Yi-Hsuan Lee, Dr. Margaret Ho, Dr. Teng-Wei Huang, Dr. Chen-Hsin Albert Yu, and the members of the Wu laboratory for helpful discussions. The authors are grateful to the National RNAi Core Facility at Academia Sinica (Taipei, Taiwan) for AAV5 packaging used in CRISPR–Cas9–mediated gene cKO; to the Imaging Core Facilities at the Institute of Molecular Biology (IMB), Academia Sinica, for technical support; and to the Bioinformatics Core Facility at IMB, Academia Sinica, for substantial assistance with analysis of the publicly available MOp dataset. The authors thank Academia Sinica Advanced Optics Microscope Core Facility for microscope imaging technical support. The core facility is funded by Academia Sinica Core Facility and Innovative Instrument Project (AS-CFII-114-A3). This work was supported by the Academia Sinica Career Development Award (AS-CDA-110-L05 to Y.-W.W.); the National Science and Technology Council, Taiwan (NSTC 113-2320-B-001-023-MY3 to Y.-W.W.; NSTC 112-2321-B-001-007 to Y.-W.W.); and start-up funds and SPP (2023) from the Institute of Molecular Biology, Academia Sinica.

## Author contributions

S.B. and Y.-W.W. designed the experiments. S.B. performed the 2-photon imaging in the acute brain slices, SIM imaging in fixed brain slices and the immunohistochemistry experiments. S.B. and M.-X.P. performed analysis of SIM imaging. K.-H.Y, C.-H.A.Y, S.B. and Y.-W.W. analyzed the publicly available scRNA-seq dataset. S.B., S.-K.T., M.-Y.C., S.B., Z.-H.Z. and S.-J.C performed experiments for confirming gene cKO. S.B., Y.-M.H, and S.-L.C. performed AAV delivery in mice. S.B., P.-Y.W. and T.-H.L. and Z.-B.T. maintained and provided transgenic mice. Y.-W.W. and S.B. wrote the manuscript with contribution of S.-J.C., S.-L.C., M.-Y.C, and all coauthors.

## STAR Methods

### Key resources table

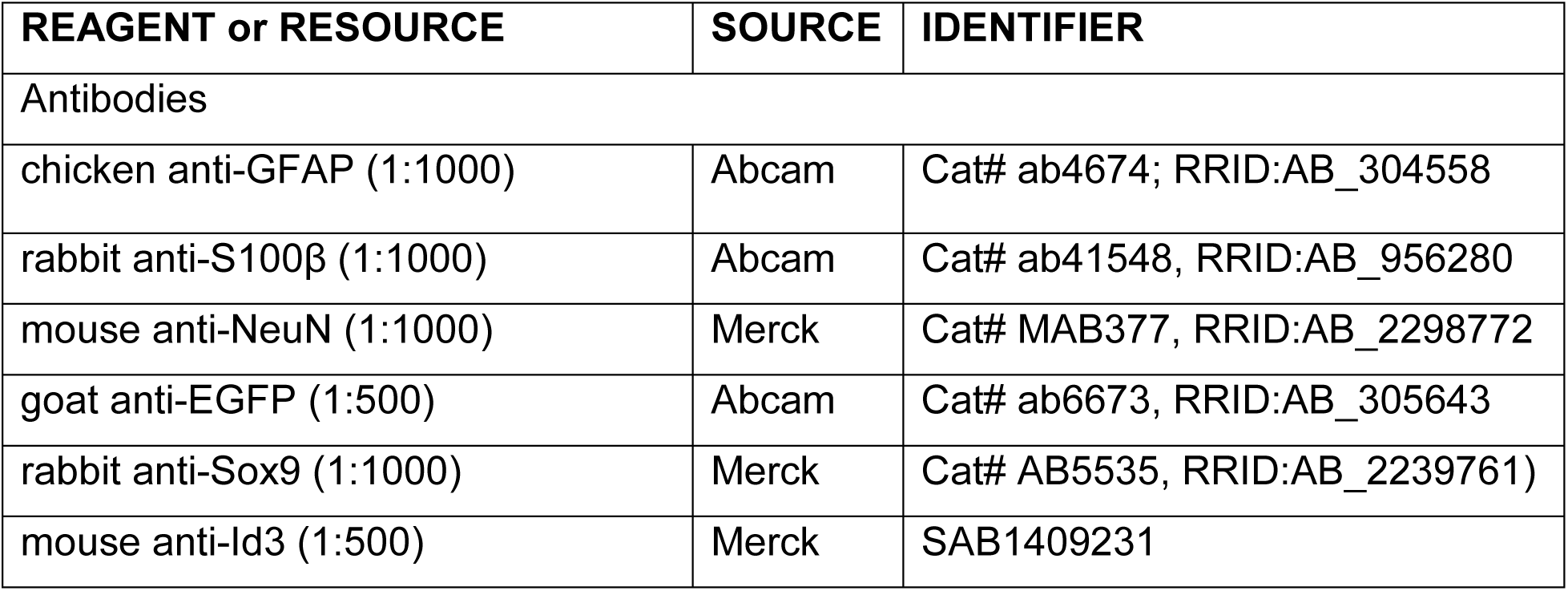
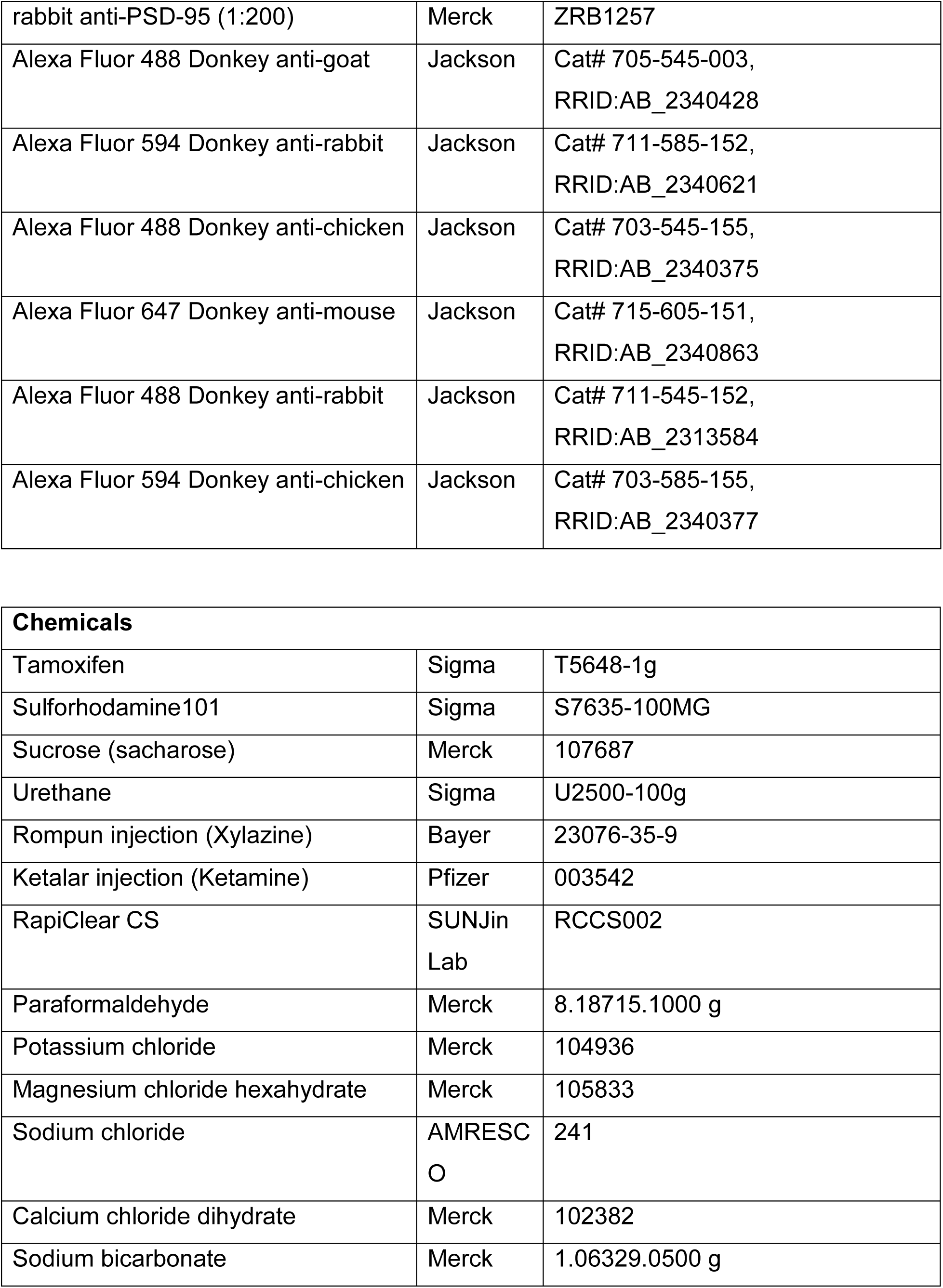
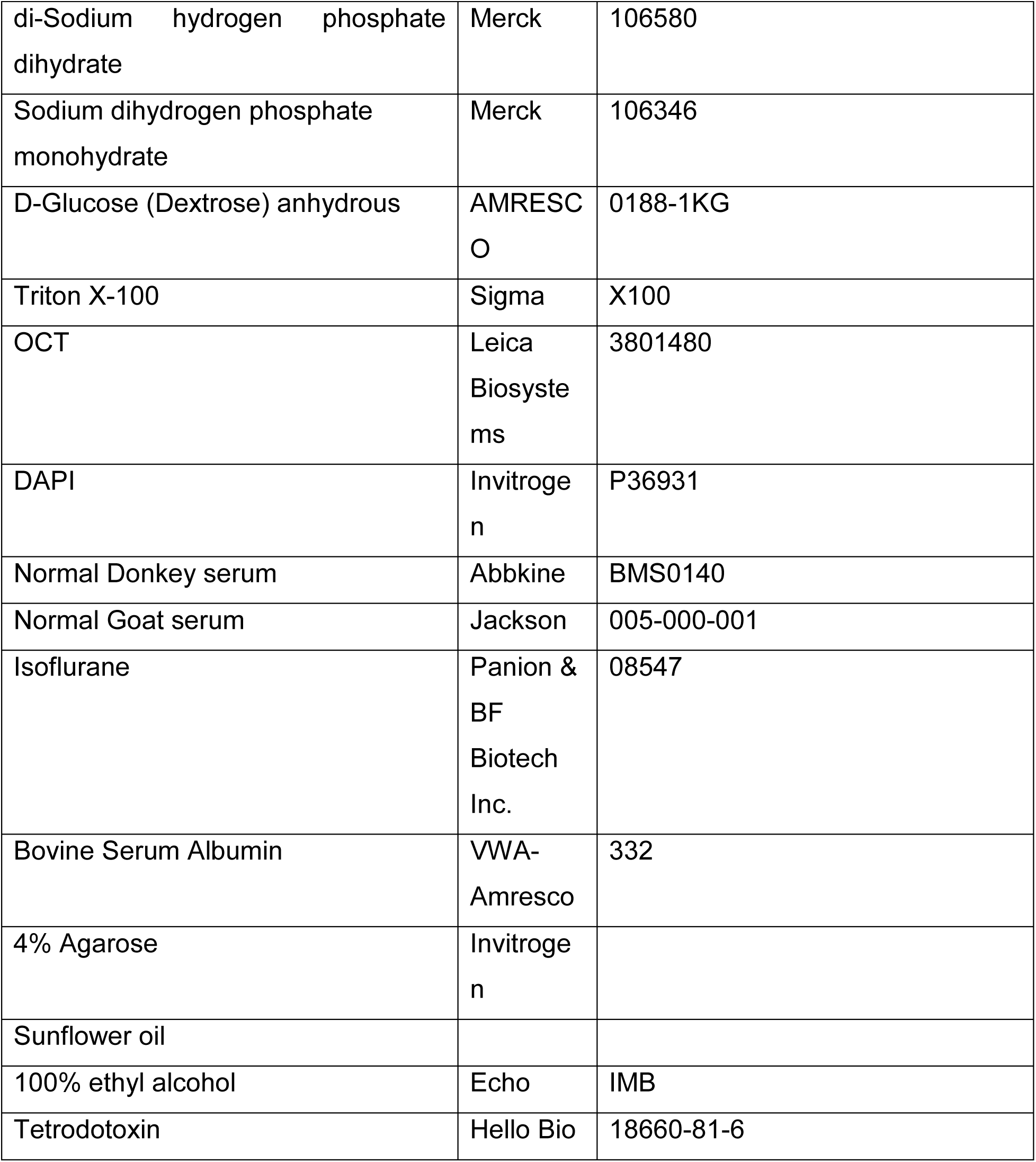

| REAGENT or RESOURCE | SOURCE | IDENTIFIER |
| --- | --- | --- |
| Antibodies |  |  |
| chicken anti-GFAP (1:1000) | Abcam | Cat# ab4674; RRID:AB_304558 |
| rabbit anti-S100β (1:1000) | Abcam | Cat# ab41548, RRID:AB_956280 |
| mouse anti-NeuN (1:1000) | Merck | Cat# MAB377, RRID:AB_2298772 |
| goat anti-EGFP (1:500) | Abcam | Cat# ab6673, RRID:AB_305643 |
| rabbit anti-Sox9 (1:1000) | Merck | Cat# AB5535, RRID:AB_2239761) |
| mouse anti-Id3 (1:500) | Merck | SAB1409231 |
| rabbit anti-PSD-95 (1:200) | Merck | ZRB1257 |
| Alexa Fluor 488 Donkey anti-goat | Jackson | Cat# 705-545-003,<br>RRID:AB_2340428 |
| Alexa Fluor 594 Donkey anti-rabbit | Jackson | Cat# 711-585-152,<br>RRID:AB_2340621 |
| Alexa Fluor 488 Donkey anti-chicken | Jackson | Cat# 703-545-155,<br>RRID:AB_2340375 |
| Alexa Fluor 647 Donkey anti-mouse | Jackson | Cat# 715-605-151,<br>RRID:AB_2340863 |
| Alexa Fluor 488 Donkey anti-rabbit | Jackson | Cat# 711-545-152,<br>RRID:AB_2313584 |
| Alexa Fluor 594 Donkey anti-chicken | Jackson | Cat# 703-585-155,<br>RRID:AB_2340377 |

| **Chemicals** |  |  |
| --- | --- | --- |
| Tamoxifen | Sigma | T5648-1g |
| Sulforhodamine101 | Sigma | S7635-100MG |
| Sucrose (sacharose) | Merck | 107687 |
| Urethane | Sigma | U2500-100g |
| Rompun injection (Xylazine) | Bayer | 23076-35-9 |
| Ketalar injection (Ketamine) | Pfizer | 003542 |
| RapiClear CS | SUNJin<br>Lab | RCCS002 |
| Paraformaldehyde | Merck | 8.18715.1000 g |
| Potassium chloride | Merck | 104936 |
| Magnesium chloride hexahydrate | Merck | 105833 |
| Sodium chloride | AMRESC<br>O | 241 |
| Calcium chloride dihydrate | Merck | 102382 |
| Sodium bicarbonate | Merck | 1.06329.0500 g |
| di-Sodium hydrogen phosphate dihydrate | Merck | 106580 |
| Sodium dihydrogen phosphate monohydrate | Merck | 106346 |
| D-Glucose (Dextrose) anhydrous | AMRESCO | 0188-1KG |
| Triton X-100 | Sigma | X100 |
| OCT | Leica Biosystems | 3801480 |
| DAPI | Invitrogen | P36931 |
| Normal Donkey serum | Abbkine | BMS0140 |
| Normal Goat serum | Jackson | 005-000-001 |
| Isoflurane | Panion & BF Biotech Inc. | 08547 |
| Bovine Serum Albumin | VWA-Amresco | 332 |
| 4% Agarose | Invitrogen |  |
| Sunflower oil |  |  |
| 100% ethyl alcohol | Echo | IMB |
| Tetrodotoxin | Hello Bio | 18660-81-6 |

#### Viral strains

|  |  |  |
| --- | --- | --- |
| AAV5 *GfaABC1D-Lck-jRGECO1a-Id1* sgRNA2&3 | This paper<br>(AS RNAi<br>Core) | NA |
| AAV5 *GfaABC1D-Lck-jRGECO1a-Id3*_sgRNA1&3 | This paper<br>(AS RNAi<br>Core) | NA |

#### Experimental model

| MOUSE LINES | SOURCE | IDENTIFIER |
| --- | --- | --- |
| C57BL/6J | Jackson<br>Laboratories | RRID:IMSR_JAX:000664<br>https://www.jax.org/strain/000664 |
| B6;FVB-Tg(Aldh1l1-cre/ERT2)1Khakh/J | Jackson<br>Laboratories | RRID:IMSR_JAX:029655<br>https://www.jax.org/strain/029655 |
| C57BL/6N-Gt(ROSA)26Sor<sup>tm1</sup>(CAG-GCaMP6f)Khakh/J | Jackson<br>Laboratories | RRID:IMSR_JAX:029626<br>https://www.jax.org/strain/029626 |
| B6.Cg-Gt(ROSA)26Sor<sup>tm9</sup>(CAG-tdTomato)Hze/J | Jackson<br>Laboratories | RRID:IMSR_JAX:007909<br>https://www.jax.org/strain/007909 |
| B6(129S4)-Gt(ROSA)26Sor<sup>tm1.1</sup>(CAG-tdTomato/GCaMP6f)Mdcah/J | Jackson<br>Laboratories | RRID:IMSR_JAX:031968<br>https://www.jax.org/strain/031968 |
| B6J.129(B6N)-Gt(ROSA)26Sor<sup>tm1</sup>(CAG-cas9*,-EGFP)Fezh/J | Jackson<br>Laboratories | RRID:IMSR_JAX:026175<br>https://www.jax.org/strain/026175 |

#### Software and Algorithms

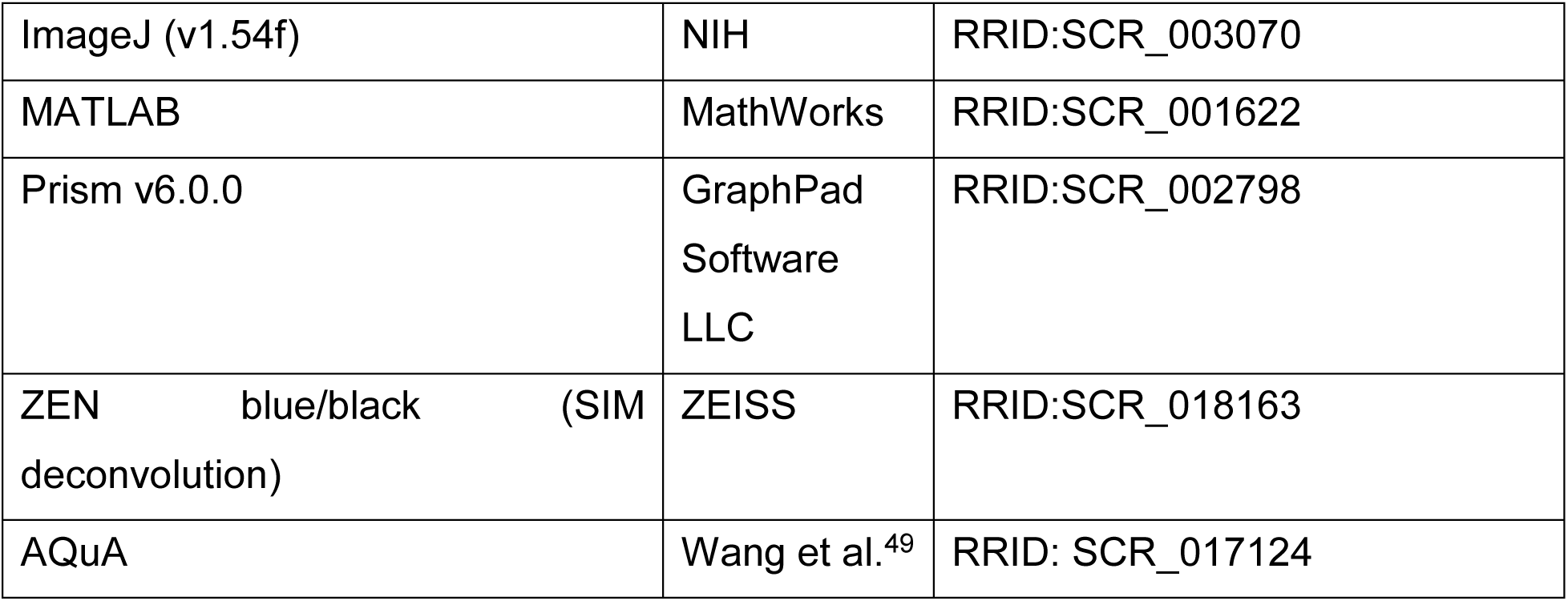

### RESOURCE AVAILABILITY

#### Lead contact

Further information and requests for resources and reagents should be directed to and will be fulfilled by the lead Contact, Dr. Yu-Wei Wu.

Dr. Yu-Wei Wu Institute of Molecular Biology, Academia Sinica, Taipei 115, Taiwan +886-2-2789-9334 (Tel)

#### Material availability statement

All materials generated in this study are available from the corresponding author upon reasonable request. Any unique reagents, data, or protocols developed for this study will be made available under standard material transfer agreements (MTAs). Publicly available datasets used in this study can be accessed as indicated in the Methods section.

#### Data and code availability

The datasets generated and analyzed during the current study are available from the corresponding author upon reasonable request. All code used for data analysis is also available upon request and will be shared under a standard open-source license. Publicly available datasets referenced in this study can be accessed as described in the Methods section. Any additional information required to reproduce the study’s findings is available from the corresponding author upon reasonable request.

### EXPERIMENTAL MODEL AND SUBJECT DETAILS

All animal experiments were conducted according to protocols approved by the Academia Sinica Institutional Animal Care & Utilization Committee. All mice were generated in the C57BL/6J genetic background. Both sexes of mice were used in experiments. Mice were maintained in 12-h light/12-h dark cycle sterile ventilated cages with access to food and water *ad libitum* at IMB animal facilities.

#### Mouse models

Experiments involved mice aged between 7 and 15 weeks, encompassing both genders. For astrocyte calcium imaging, heterozygous Aldh1l1-CreER^T2^ mice (JAX Stock No. 029655)^48^ were crossbred with homozygous Cre-dependent Lck-GCaMP6f knock-in mice (JAX Stock No. 029626)^48^. This breeding produced Aldh1l1-CreER^T2^; Lck-GCaMP6f double transgenic mice for detailed calcium imaging studies. In parallel, for morphological analysis, heterozygous Aldh1l1-CreER^T2^ mice were mated with homozygous Rosa-CAG-LSL-tdTomato (Ai9, JAX Stock No.007909)^77^ mice. For astrocyte-specific gene knockout experiments, the strategy involved breeding heterozygous Aldh1l1-CreER^T2^ mice with homozygous Cre-dependent Rosa26-LSL-Cas9-EGFP knock-in mice (LSL-Cas9-EGFP; JAX Stock No. 026175)^70^ to obtain Aldh1l1-CreER^T2^;LSL-Cas9-EGFP double transgenic mice.

### Method details

#### Tamoxifen induction of CreER^T2^-dependent gene expression

Injectable tamoxifen (Sigma #T5648) was freshly prepared by combining a tamoxifen stock solution (100 mg/ml) with sunflower oil at a 1:9 ratio. This formulation was used for sparsely labeling astrocytes with fluorescent reporters in various transgenic mouse models: Aldh1l1-CreER^T2^;Lck-GCaMP6f, Aldh1l1-CreER^T2^;LSL-tdTomato mice. A single intraperitoneal (*i.p.*) injection of tamoxifen at a dose of 50 mg/kg body weight was administered to 4-week-old mice for this purpose. These mice were used for further imaging and recording three weeks post induction. For inducible gene knockout experiments, Aldh1l1-CreER^T2^;LSL-Cas9-EGFP mice, injected with AAV5 GfaABC1D-Lck-jRGECO1a-Mus Id1 sgRNA2&3 and AAV5 GfaABC1D-Lck-jRGECO1a-Mus Id3 sgRNA1&3 viruses at 4-5 weeks age, were induced at 8-9 weeks of age. Five consecutive days injection of tamoxifen at a dose of 100 mg/kg body weight was administered to achieve sparse astrocyte-specific KO. These mice were used for further imaging and recording one-month post induction. These protocols were designed to selectively manipulate gene expression in astrocytes, enabling the study of their specific roles in various genetic contexts.

The tamoxifen stock solution was prepared by dissolving 20 mg tamoxifen powder in 500 µl of 100% ethyl alcohol. This was facilitated using a sonicator with controlled heating at 40-50°C for 15-30 minutes. The prepared stock was then added into 980 µl of sunflower oil31. This solution was then further dissolved in a drying vacuum oven for 15-20 minutes, until the volume reduced to 1000 µl. This is then stored at -20°C and used within 8 weeks to ensure potency.

#### Acute brain slice preparation

Mice aged between 7 to 15 weeks (at least 3 weeks post tamoxifen-induction) were first administered with Sulforhodamine 101 (SR101) dye (5mg/ml in 0.9% saline) intraperitoneally (*i.p.*), one hour prior to decapitation^47,78,79^. SR101, a red fluorescent dye, selectively stains astrocyte soma and their thicker processes. Following deep anesthesia with isoflurane (no toe reflex detected), mice were decapitated, and their brains were immediately extracted and immersed in ice-cold slush of artificial cerebrospinal fluid (ACSF). Coronal brain sections, 300 µm in thickness, were prepared using a vibratome (Leica VT1200 S, Germany) at a speed of 0.14 mm/s. The ACSF used for slicing contained the following components (in mM): NaCl 125, KCl 2.5, Na_2_HPO_4_.H_2_O 1.25, NaHCO_3_ 25, D-glucose 15, CaCl_2_.2H_2_O 2, MgCl_2_.6H_2_O 1. This solution was continuously saturated with a gas mixture of 95% O_2_ and 5% CO_2_. After slicing, the brain slices were incubated in bubbling ACSF at 34-35 °C for 30 minutes, followed by a subsequent 30-minute incubation at room temperature to allow for adequate recovery. Imaging experiments were conducted at a controlled temperature of 30°C, with brain slices continuously perfused with ACSF and constantly saturated with carbogen (95% O_2_ and 5% CO_2_). This setup ensured optimal conditions for observing astrocytic activity and morphology under the microscope. The imaging was conducted within 5 hours to ensure recording from healthy, viable cells.

#### Two-photon microscopy for acute brain slices

##### Astrocyte intracellular calcium imaging

The acute brain slices were secured in the perfusion chamber under the objective using a harp. The slices were imaged at the MOp on both the hemispheres. The location of the individual layers of the MOp was confirmed with IR-scanning differential interference contrast (DIC) imaging. We ensured that the astrocytes chosen for recording were healthy based on whether they show no clumped processes in their territory volume. Lck-GCaMP6f calcium activity was recorded under the 2P microscope (FemtoSmart Dual, Femtonics, Hungary) using a 20x/NA 1.0 objective (XLUMPLFLN Objective, Olympus, Japan) with a Ti-sapphire femtosecond laser tuned to 920 nm (Chameleon Ultra II, Coherent, USA). The laser intensity was usually 10-15% with an output of 25-30 mW and frame size of 85 μm-by-85 μm. Spontaneous calcium signals from single astrocyte were imaged from five planes (z-interval = 30 μm) using a resonant scanner with a frame rate of 31 Hz and a volume rate of 6.18 Hz. The SR101 occupying the red channel was used to identify the mid plane of the astrocyte that includes the soma.

##### Astrocyte morphology imaging

Morphology imaging was recorded using FemtoSmart Dual 2P microscope equipped with a 20x, NA1.0 objective with high power femtosecond laser (Fidelity-2, Coherent, USA) at 1070 nm. The laser intensity was usually 5% with an output of 30 mW and frame size of 85 μm-by-85 μm. The astrocytes were imaged for its entire volume with a z-step of 0.5 µm and a resolution of 1024 x 1024 pixels under the galvo mode. The laser intensity was increased with the depth of imaging. Astrocytes with their somas in close proximity were excluded from analysis. Cells occupying independent territories without any overlap with neighboring cells were chosen for imaging and analysis.

##### Astrocyte with CRISPR-Cas9 mediated knockout imaging of intracellular calcium and morphology

The acute brain slices were secured in the perfusion chamber under the objective using a harp. The slices were imaged at the MOp on both the hemispheres. The location of the individual layers of the MOp was confirmed with IR-scanning differential interference contrast (DIC) imaging. Lck-jRGECO1a calcium activity was recorded under the 2P microscope using a 20x/NA 1.0 objective with a femtosecond laser at 1070 nm (Fidelity-2, Coherent, USA). The laser intensity was usually 25-30% with an output of 140-150 mW and frame size of 85 μm-by-85 μm. Spontaneous calcium signals from single astrocyte were imaged from six planes (z-interval = 30 μm) using a resonant scanner with a frame rate of 31 Hz and a volume rate of 5.15 Hz.

The same cell is imaged in the green channel using a Ti-sapphire femtosecond laser tuned to 920 nm (Chameleon Ultra II, Coherent, USA), to confirm its expression of Cas9-EGFP. The laser intensity was usually 13-15% with an output of 30-45 mW and frame size of 85 μm-by-85 μm. The astrocytes were imaged for its entire volume with a z-step of 0.5 µm and a resolution of 512 x 512 pixels under the resonant mode, using 100 planes for averaging. The laser intensity was increased with the depth of imaging. The modes were switched in either way when exploring for the presence of knockout cells in acute brain slices.

##### Post hoc imaging of acute brain slices

The correlation of calcium activity to GFAP expression levels in astrocytes was achieved by fixing the acute brain slices imaged, with 4% paraformaldehyde. These slices were incubated for 24 hours with primary antibodies against GFAP and eGFP, following the immunohistochemistry protocol, mentioned in details later. The cells recorded for their calcium activities were identified post hoc in the fixed slices and their GFAP expression levels were analyzed.

#### Pharmacology study

Spontaneous calcium activity from each cell was initially recorded for five minutes. This was followed by the directly addition of 50 µl of 1 mM tetrodotoxin (TTX; Hello Bio. UK) stock solution into the perfusion solution of 50 ml ACSF to yield a final concentration of 1 µM. The TTX was allowed to completely mixed in the ACSF for 3 minutes. The same cells were then recorded in the presence of 1 µM TTX to study its effect on the calcium activity.

#### AAV cloning and packaging

The astrocyte-specific knockout of *Id1* and *Id3* genes in Aldh1l1-CreER^T2^;LSL-Cas9-EGFP double transgenic mice was achieved by two pairs of single-strand guide RNAs (sgRNAs) against the complementary DNA sequence (CDS) for each gene. The efficiency of these sgRNA pairs were determined by reporter assay and the pair with higher efficiency of knockout was chosen for further construction of the AAV5 virus constructs. To monitor the calcium activity in *Id1* and *Id3* KO astrocytes, we incorporated *GfaABC1D-Lck-jRGECO1a*, a membrane-tethered red-shifted genetically-encoded calcium sensor, with the sgRNAs in the same AAV plasmid. First, the pAAV.GfaABC1D-Lck-jRGECO1a plasmid was constructed using plasmid pAAV.Syn.NES-jRGECO1a.WPRE.SV40 (https://www.addgene.org/100854/) as the backbone. The *GfaABC1D* promoter was obtained from the plasmid pZac2.1 GfaABC1D-NAPA-A (https://www.addgene.org/92281/) and the *Lck-jRGECO1a* gene sequence containing plasmid was obtained from Dr. Ching-Lung Hsu’s laboratory at the Institute of Biomedical Sciences of Academia Sinica. The sgRNA cassettes were then inserted into the construct. The final constructs after packaging into AAV5 are (1) AAV5-GfaABC1D-Lck-jRGECO1a-Id1 sgRNA2&3 and (2) AAV5-GfaABC1D-Lck-jRGECO1a-Id3_sgRNA1&3.

#### Stereotaxic microinjections of AAVs

Double transgenic Aldh1l1-CreER^T2^;LSL-Cas9-EGFP mice are anesthetized and placed onto a stereotaxic frame. Continuous anesthesia using isoflurane (1.5%) was carefully monitored and maintained throughout the surgery. The skin covering the head was cut to expose the skull. The area was sterilized using povidone and 75% ethanol. Glass pipettes (World Precision Instruments) were pulled to a tip diameter of 30-35 μm with an angle of 30° and filled with the desired virus. The tip diameter and angle are optimized to minimize the extent of invasion. The viruses were injected at two depths from the pial surface (0.4 mm and 0.6 mm). A total volume of 1000 nl was injected at each of the above depths of the primary motor cortex (AP 1.0 mm and ML-1.0 mm) unilaterally. The glass pipettes are inserted to the desired depth and kept there for 10 minutes. AAVs were injected at a rate of 1 nl/sec using Nanoject III injector (Drummond SCI). Glass pipettes are withdrawn 10 min after injection is complete and scalps are cleaned and sutured with sterile surgical silk sutures (UNIK Sutures, SC125). Mice were allowed to recover in clean cages. Viruses used in this study are AAV5-GfaABC1D-Lck-jRGECO1a-Mus_Id1_sgRNA2_3 and AAV5-GfaABC1D-Lck-jRGECO1a-Mus_Id3_sgRNA1_3. Equal volumes of these viruses were mixed and then injected at the desired depths.

#### Cardiac perfusion

Mice were put in a chamber containing isoflurane for deep anesthesia. This was followed by *i.p.* injection with urethane (1.5 g/kg). Once all the reflexes subsided, the abdominal cavity was opened up. A butterfly needle was pierced into the left ventricle of the heart and a cut was made in the right atrium precisely, to ensure the lungs are not punctured. The mice were perfused with 50 ml of 0.9% freshly prepared cold saline until all the blood flows out. This was followed by perfusion with 25 ml of ice cold 4% paraformaldehyde (PFA). A successful perfusion can be ensured by the body twitching of the mice upon PFA perfusion. The mice body will become hard, a key indicator being the liver which also turns harder. The head was slit and the skull was peeled carefully so as to extract the brain. A white brain with no traces of blood is a good indicator for a successful perfusion. The brains were post fixed overnight (16 hours) at 4°C, in 4% PFA under shaking conditions (120 rpm/min). The following day, the brains were transferred into cold 30% sucrose solution in 0.1M PB for at least 48 hours (with constant agitation at 4°C) until further use. The brains should sink at the bottom of the vial.

#### Immunohistochemistry (IHC)

Mice brains were embedded in OCT and coronal sections of the brain (50 µm) were obtained using Leica’s cryostat microtome (-20°C) and collected in 0.1 M PB. The slices were then washed thrice in 0.01 M PBS for 10 minutes each at RT (with agitation, 120 rpm/min). The slices were then blocked for 1 hour in freshly prepared 2% normal donkey serum (NDS), 10% bovine serum albumin (BSA) dissolved in 0.5% Triton X-100 in 0.01 M PBS at RT, with agitation. The antibodies added were as follows: (1) chicken anti-GFAP (1:1000), rabbit anti-S100β (1:1000), mouse anti-NeuN (1:1000), (2) chicken anti-GFAP (1:1000), rabbit anti-Sox9 (1:1000), mouse anti-NeuN (1:1000), (3) goat anti-EGFP (1:500), chicken anti-GFAP (1:1000), mouse anti-NeuN (1:1000), (4) chicken anti-GFAP (1:1000), rabbit anti-Sox9 (1:1000), mouse anti-Id3 (1:500). These antibodies were diluted in the above blocking solution and incubated with the brain slices for O/N at 4°C with agitation at 120 rpm/min. The following day, the slices were washed thrice in 0.01 M PBS for 10 minutes each. The secondary antibodies used are as follows: Alexa Fluor 488 Donkey anti-goat, Alexa Fluor 594 Donkey anti-rabbit, Alexa Fluor 488 Donkey anti-chicken, Alexa Fluor 647 Donkey anti-mouse, Alexa Fluor 647 Donkey anti-rabbit, Alexa Fluor 488 Donkey anti-rabbit, Alexa Fluor 594 Donkey anti-chicken, along with DAPI (1:5000). These antibodies were diluted in the above blocking solution and incubated with the brain slices for 2 hours at 4°C with agitation at 120 rpm/min. The slices were then washed thrice in 0.01 M PBS for 10 minutes each. Following washing, the slices were mounted on slides. Upon drying, clearing agent, RapiClear CS (https://www.sunjinlab.com/product-category/rapiclear/) was added to each slice on the slide and covered with coverslip. The images were taken using 20X/0.8 Plan Apochromat objective lens in Zeiss LSM710 and LSM780 laser scanning microscopes.

#### Structured illumination microscopy (SIM)

Coronal sections (20-25 µm in thickness) of Aldh1l1-CreER^T2^; LSL-tdTomato mice brain were obtained using Leica’s cryostat microtome (-20°C) and collected in 0.1 M PB. The sections were then washed thrice in 0.01 M PBS for 10 minutes each at RT (with agitation at 120 rpm/min). The slices were incubated with DAPI (1:5000) for 2 hours at 4°C (with agitation at 120 rpm/min) followed by washing thrice in 0.01 M PBS at RT. DAPI helps in identification of the layers the astrocytes belong to. The slices were then mounted on slides (one per slide) using RapiClear CS and covered with coverslips (Zeiss High-Performance). The sections were stored at 4°C until use. Imaging was performed using 63X/1.4 Plan Apochromat objective lens with 561 nm laser of Zeiss Elyra SP1 microscope or Zeiss ELYRA 7 with a grating set to 5. With Zeiss ELYRA 7 microscope and its improved SIM^2^ processing algorithm, the lateral resolution of approaches ∼60 nm, whereas the axial resolution is improved to ∼200 nm. A Z-step of 126 nm was taken for astrocytes imaged from different regions. A laser power of 5% was used for lasers 488 nm and 561 nm. From each astrocyte, three regions of interest were picked and they were further analyzed for specialized loop-like structures.

#### PSD-95 co-localization with LLS

Aldh1l1-CreER^T2^;LSL-tdTomato mice brain coronal sections (25 µm in thickness) were obtained using Leica’s cryostat microtome (-20°C) and collected in 0.1 M PB. The slices were then washed thrice in 0.01 M PBS for 10 minutes each at RT (with agitation, 120 rpm/min). The slices were then blocked for 1 hour in freshly prepared 2% normal donkey serum (NDS), 10% bovine serum albumin (BSA) dissolved in 0.5% Triton X-100 in 0.01 M PBS at RT, with agitation. The slices were then incubated with rabbit anti-PSD-95 (1:400) O/N with agitation at 4°C. The following day, the slices were washed thrice in 0.01 M PBS for 10 minutes each. The slices were incubated with DAPI (1:5000) and secondary antibody Alexa Fluor 488 Donkey anti-rabbit for 2 hours at 4°C (with agitation at 120 rpm/min) followed by washing thrice in 0.01 M PBS at RT. DAPI helps in identification of the layers the astrocytes belong to. The slices were then mounted on slides (one per slide) using RapiClear CS and covered with coverslips (Zeiss High-Performance). The sections were stored at 4°C until use. Imaging was performed using 63X/1.4 Plan Apochromat objective lens with 488 nm and 561 nm laser of Zeiss Elyra SP1 microscope with a grating set to 5. A Z-step of 126 nm was taken for astrocytes imaged from different regions. From each astrocyte, three regions of interest were picked and they were further analyzed for specialized loop-like structures.

#### Imaging analysis for calcium imaging using AQuA

The raw recording was opened up using ImageJ. A walking average of 2 frames was applied on each of the recordings. They were then registered using custom written MATLAB code. The recordings were then analyzed using AQuA. The spatial and temporal resolutions of our recordings were as follows: 0.166218 µm per pixel and 6.18 Hz. We specify the astrocyte territory by a manually drawn ROI. We removed pixels whose distance was smaller or equal to 5 pixels from the border. In the SIGNAL TAB, we applied a thrARScl of 3, smoXY as 1 and minSize as 180-200 pixels. Following that in the VOXEL TAB, we applied a thrTWScl to 2, thrExtZ to 1 or 2, depending on the signal. In the EVENT TAB, we chose the following parameters: cRise=2 cDelay=2 and gtwSmo=1. Lastly, we applied a zThr to 2.

#### Imaging analysis for SIM imaging

The raw images were first processed using SIM^2^ algorithm and then opened up using ImageJ. An ROI of area 57.973 μm^2^ was created. A total of 3 ROIs were chosen from each image Z-stack. It was made sure that the ROIs are in the fine processes. The same ROI size was used across all astrocytes recorded from different regions. The ROIs were then dug in to look for LLSs. A loop-like structure is defined by its presence for at least 3 frames (∼378 nm). We classified them into different types based on their shapes. The total number of loop-like structures, their inner area and perimeter were quantified in order to compare across regions.

In case of the PSD-95 and LLS co-localization analysis, the channels were separated into two and the LLSs were quantified first. These LLSs are then analyzed further for the co-localization with PSD-95 by merging the channels together. This allows an unbiased way to calculate the percentage of LLSs that co-localize with PSD-95. The random shuffling was performed by merging tdTomato channel of one ROI with the PSD-95 channel of another ROI and vice versa (**Figure S2E**).

#### Single cell-RNA-seq data analysis

The Brain Initiative Cell Census Network (BICCN) generated 10X Chromium dataset (Broad) of scRNA-sequencing of the primary motor cortex (MOp)^1,52,80^ is used in this study. Focusing primarily on the astrocytes, with the help of our Bioinformatics core in IMB, we generated the UMAP featuring a total of 18 clusters using a resolution of 1.0. The genes are plotted in a heatmap that includes the top 10 genes of each cluster (**S8C**).

The L1 cluster confirmation was performed using the MERFISH dataset, MERSCOPE Dataset: Michael Kunst, Delissa McMillen, Jennie Close, Jazmin Campos, Madie Hupp, Naomi Martin, Jocelin Malone, Zoe Maltzer, Augustin Ruiz, Nasmil Valera Cuevas, Brian Long, Jack Waters, Hongkui Zeng. (2023). Whole Mouse Brain Transcriptomic Cell Type Atlas - MERSCOPE v1^52^.

### QUANTIFICATION AND STATISTICAL ANALYSIS

The sample sizes, statistical tests, statistical comparisons, n numbers and p values are shown in the figure panels or figure legends and also stated within the main text and figure legends. Throughout the manuscript, N is defined as the numbers of mice used in this study while definition of n is case-dependent, either numbers of astrocytes or brain slices; stated in the respective figure legends. Statistical tests were run in GraphPad Prism 6. Summary data are presented as mean ± SEM along with the individual data points. The details of the mean and SEM values are provided in the main text or supplementary figure legends. We employed non-parametric tests due to the sample sizes and non-Gaussian distribution of the data. In the figures, p values are indicated by asterisk(s): *p < 0.05; **p < 0.01; ***p < 0.001; ****p < 0.0001.

