## Supplementary figures and legends for "A Distinct Layer 1 Astrocyte Program Shapes Perisynaptic Structure and Calcium Signaling in Mouse Motor Cortex"

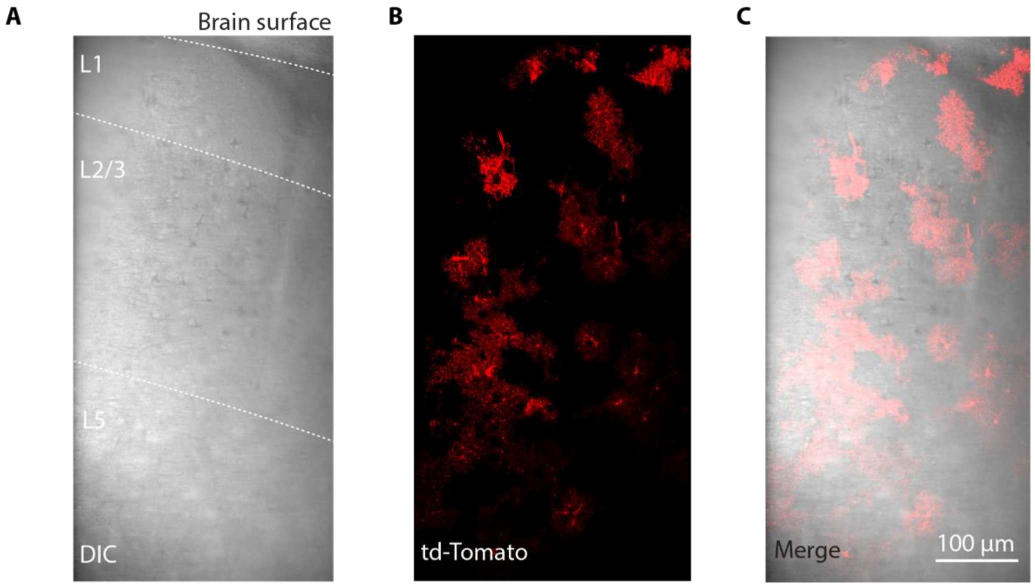


**Figure S1. IR-DIC imaging used as a reference to demarcate the layers of the MOp.**

(A) IR-DIC imaging used for demarcation of MOp layers. The white dashed lines show the boundary of each layer.

(B) Sparsely labelled td-Tomato in astrocytes imaged across the cortex.

(C) IR-DIC image merged over td-Tomato to facilitate accurate astrocyte imaging from each layer.

Scale bar represent 100 µm.

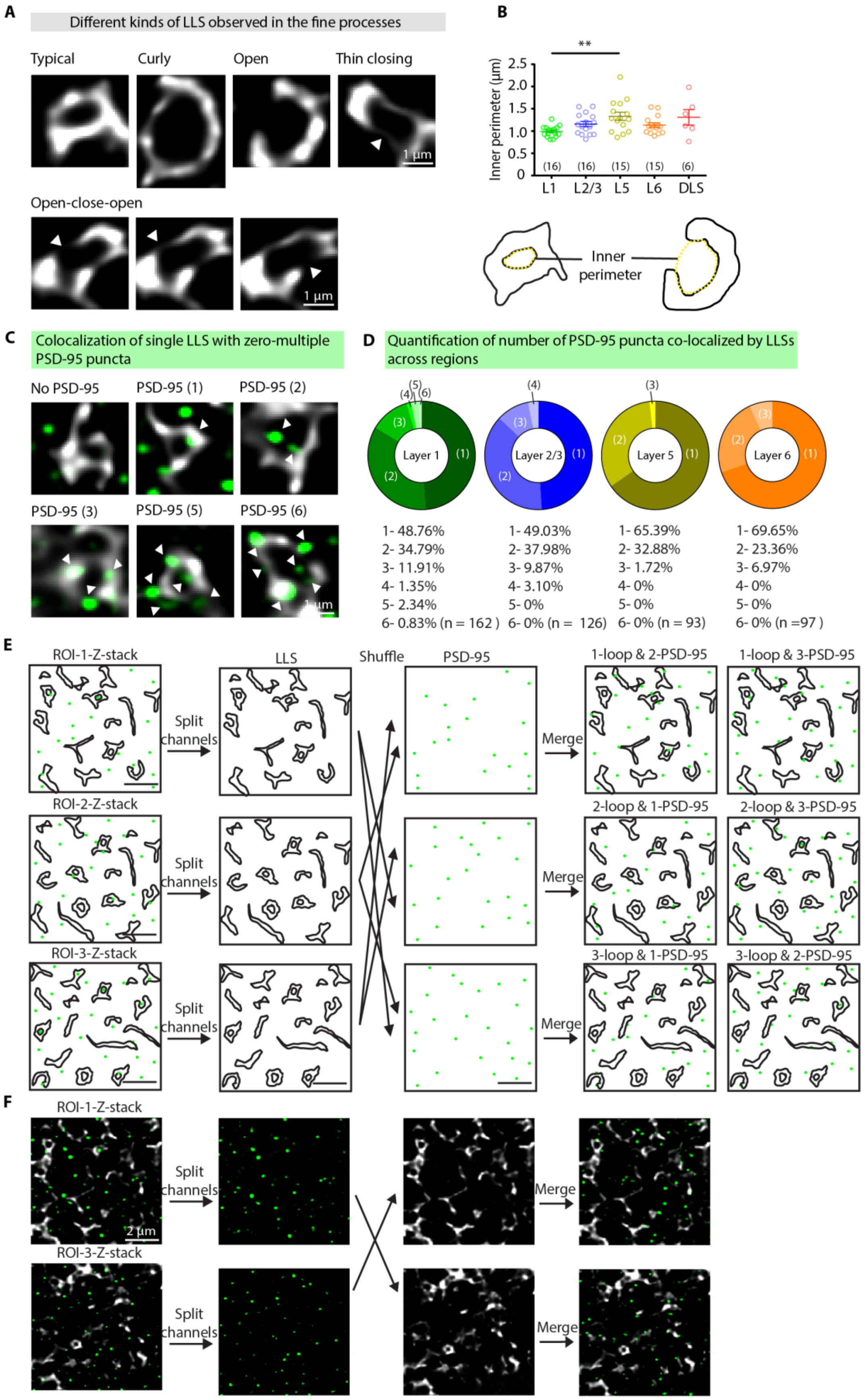
A
Different kinds of LLS observed in the fine processes
Typical
Curly
Open
Thin closing
1 µm
Open-close-open
1 µm
B
**
Inner perimeter (µm)
2.5
2.0
1.5
1.0
0.5
0
(16)
(16)
(15)
(15)
(6)
L1
L2/3
L5
L6
DLS
Inner perimeter
C
Colocalization of single LLS with zero-multiple PSD-95 puncta
No PSD-95
PSD-95 (1)
PSD-95 (2)
PSD-95 (3)
PSD-95 (5)
PSD-95 (6)
D
Quantification of number of PSD-95 puncta co-localized by LLSs across regions
Layer 1
Layer 2/3
Layer 5
Layer 6
1- 48.76%
2- 34.79%
3- 11.91%
4- 1.35%
5- 2.34%
6- 0.83% (n = 162 )
1- 49.03%
2- 37.98%
3- 9.87%
4- 3.10%
5- 0%
6- 0% (n = 126)
1- 65.39%
2- 32.88%
3- 1.72%
4- 0%
5- 0%
6- 0% (n = 93)
1- 69.65%
2- 23.36%
3- 6.97%
4- 0%
5- 0%
6- 0% (n =97 )
E
ROI-1-Z-stack
ROI-2-Z-stack
ROI-3-Z-stack
Split channels
LLS
Shuffle
PSD-95
Merge
1-loop & 2-PSD-95
1-loop & 3-PSD-95
2-loop & 1-PSD-95
2-loop & 3-PSD-95
3-loop & 1-PSD-95
3-loop & 2-PSD-95
F
ROI-1-Z-stack
ROI-3-Z-stack
Split channels
Merge
2 µm

**Figure S2. Detailed LLS morphology types and quantification across the layers of the MOp and DLS.**

(A) Examples of the different kinds of LLSs observed in astrocytes. Typical- has a typical loop like structure that lasts for at least 3 frames or 378 nm. Curly- these LLSs have an uneven structure, often curly in appearance. These either have a thin closing or do not close onto itself. Open- these LLSs do not close but have a semi-circular structure (>50%). Thin closing- these LLS have a very thin flap like closing that's comparatively thinner than the rest of the LLS. Open-close-open- these LLSs are rare, lasts for 3 frames. These frames display an open LLS, that closes in the second frame and then open immediately in the subsequent frame.

(B) (Top) Quantification of the inner perimeter of the LLS across regions. (L1: 0.99 ± 0.03 µm, n = 16 ROIs/ 6 cells; L2/3: 1.16 ± 0.06 µm, n = 16 ROIs/ 6 cells; L5: 1.33 ± 0.09 µm, n = 15 ROIs/5 cells; L6: 1.14 ± 0.05 µm, n = 15 ROIs/ 5 cells; DLS: 1.31 ± 0.17 µm, n = 6 ROIs/ 2 cells; N = 6 mice; Kruskal-Wallis test followed by Dunn's multiple comparisons test) (bottom) Illustration showing examples of LLSs with the yellow dashed line representing its inner perimeter.

(C) Examples of the LLSs in contact with multiple PSD-95 puncta.

(D) Quantification of the number of PSD-95 puncta co-localized with LLS, across regions. (n = 2 astrocytes; 6 ROIs; 162 LLSs (L1), n = 2 astrocytes; 6 ROIs; 123 LLSs (L2/3), n = 2 astrocytes; 6 ROIs; 93 LLSs (L5), n = 2 astrocytes; 6 ROIs; 97 LLSs (L6), N = 4 mice).

(E) Illustration to show pipeline used to generate the random shuffling data.

(F) Original imaging data to support the above pipeline.

The data are shown as mean ± SEM.

**$p < 0.01$.

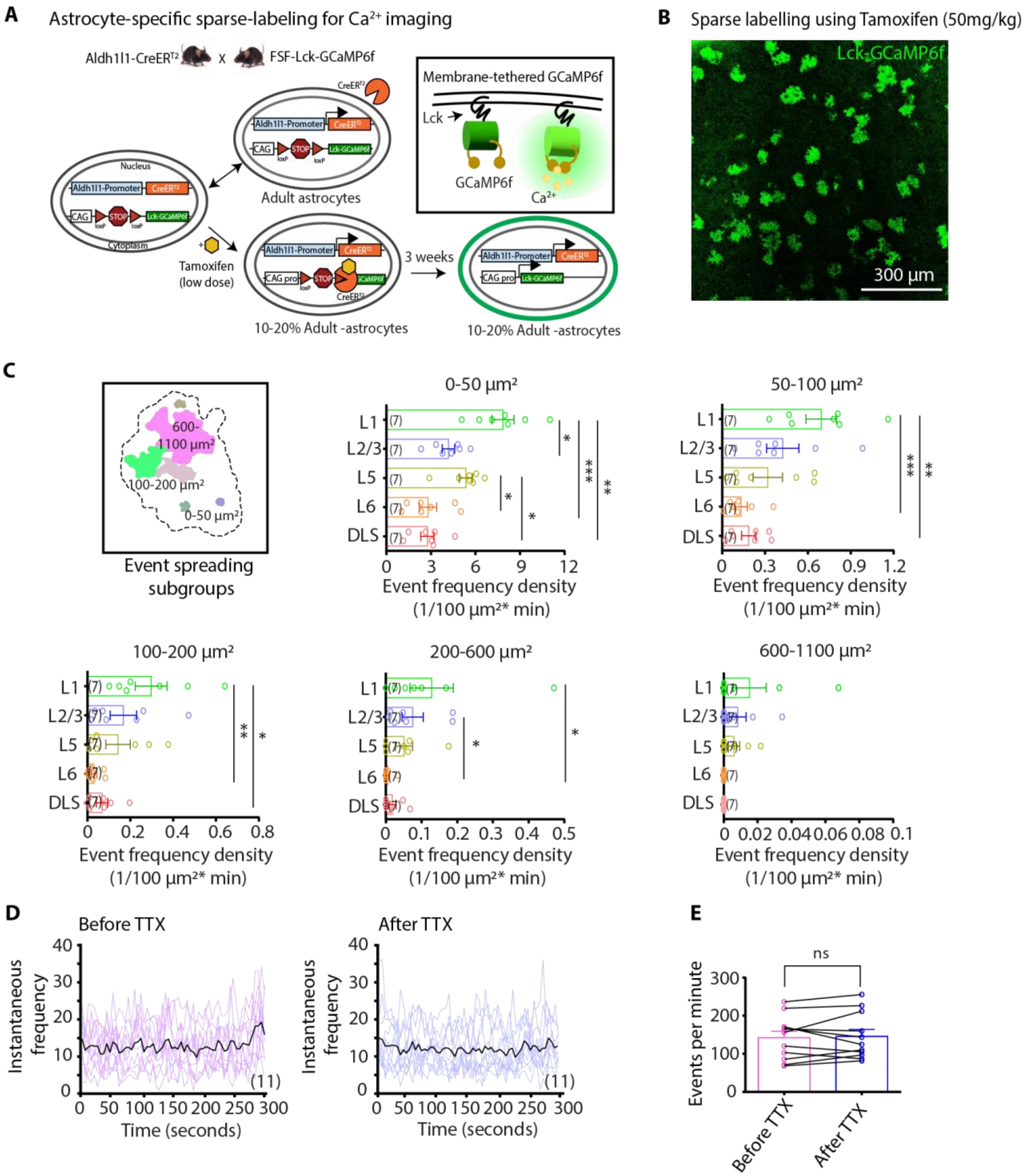


**Figure S3. Calcium event frequency density of the spreading subgroups across the MOp and the DLS.**

(A) Mechanism of Tamoxifen mediated induction of Lck-GCaMP6f gene in astrocytes.
(B) Sparse labeling of astrocytes in the MOp obtained from 50 mg/kg Tamoxifen injection.
(C) Event frequency density of spreading subgroups, quantified across regions. (n = 7 (L1), n = 7 (L2/3), n = 7 (L5), n = 7 (L6), n = 7 (DLS) astrocytes; N = 14 mice; Kruskal-Wallis test followed by Dunn's multiple comparisons test). The data are shown as mean ± SEM.

(0-50 $\mu m^2$; L1: 7.9 ± 0.7 events/$10^2$ $\mu m^2$ per minute, n = 7, L2/3: 4.2 ± 0.4 events/$10^2$ $\mu m^2$ per minute, n = 7, L5: 5.4 ± 0.5 events/$10^2$ $\mu m^2$ per minute, n = 7, L6: 2.8 ± 0.6 events/$10^2$ $\mu m^2$ per minute, n = 7, DLS: 2.8 ± 0.4 events/$10^2$ $\mu m^2$ per minute, n = 7; 50-100 $\mu m^2$; L1: 0.0069 ± 0.0010 events/$10^2$ $\mu m^2$ per minute, n = 7, L2/3: 0.0042 ± 0.0011 events/$10^2$ $\mu m^2$ per minute, n = 7, L5: 0.0032 ± 0.0010 events/$10^2$ $\mu m^2$ per minute, n = 7, L6: 0.0013 ± 4 x $10^{-4}$ events/$10^2$ $\mu m^2$ per minute, n = 7, DLS: 0.0018 ± 4 x $10^{-4}$ events/$10^2$ $\mu m^2$ per minute, n = 7; 100-200 $\mu m^2$; L1: 0.0029 ± 7 x $10^{-4}$ events/$10^2$ $\mu m^2$ per minute, n = 7, L2/3: 0.0016 ± 6 x $10^{-4}$ events/$10^2$ $\mu m^2$ per minute, n = 7, L5: 0.0014 ± 5 x $10^{-4}$ events/$10^2$ $\mu m^2$ per minute, n = 7, L6: 3 x $10^{-4}$ ± 1 x $10^{-4}$ events/$10^2$ $\mu m^2$ per minute, n = 7, DLS: 6 x $10^{-4}$ ± 2 x $10^{-4}$ events/$10^2$ $\mu m^2$ per minute, n = 7; 200-600 $\mu m^2$; L1: 0.0012 ± 6 x $10^{-4}$ events/$10^2$ $\mu m^2$ per minute, n = 7, L2/3: 7 x $10^{-4}$ ± 2 x $10^{-4}$ events/$10^2$ $\mu m^2$ per minute, n = 7, L5: 5 x $10^{-4}$± 2 x $10^{-4}$ events/$10^2$ $\mu m^2$ per minute, n = 7, L6: 5 x $10^{-5}$ ± 4 x $10^{-5}$ events/$10^2$ $\mu m^2$ per minute, n = 7, DLS: 1 x $10^{-4}$ ± 1 x $10^{-4}$ events/$10^2$ $\mu m^2$ per minute, n = 7; 600-1100 $\mu m^2$; L1: 1 x $10^{-4}$ ± 9 x $10^{-5}$ events/$10^2$ $\mu m^2$ per minute, n = 7, L2/3: 8 x $10^{-5}$ ± 4 x $10^{-5}$ events/$10^2$ $\mu m^2$ per minute, n = 7, L5: 6 x $10^{-5}$ ± 3 x $10^{-5}$ events/$10^2$ $\mu m^2$ per minute, n = 7, L6: 0 ± 0, n = 7 events/$10^2$ $\mu m^2$ per minute, DLS: 0 ± 0 events/$10^2$ $\mu m^2$ per minute, n = 7; N = 14 mice, Kruskal-Wallis test followed by Dunn's multiple comparisons test).

(D) Instantaneous event frequency before and after addition of TTX.

(E) Quantification of event frequency per minute before and after addition of TTX. (n = 7 (L1), n = 4 (L2/3) astrocytes, N = 7 mice; Wilcoxon matched-pairs signed-rank test).

Baseline: 142.1 ± 17.22 events per minute, n = 11, TTX: 145.6 ± 18.23 events per minute, n = 11, Wilcoxon matched-pairs signed-rank test.

The data are shown as mean ± SEM.

*p < 0.05, **p < 0.01, ***p < 0.001, ns = non-significant.

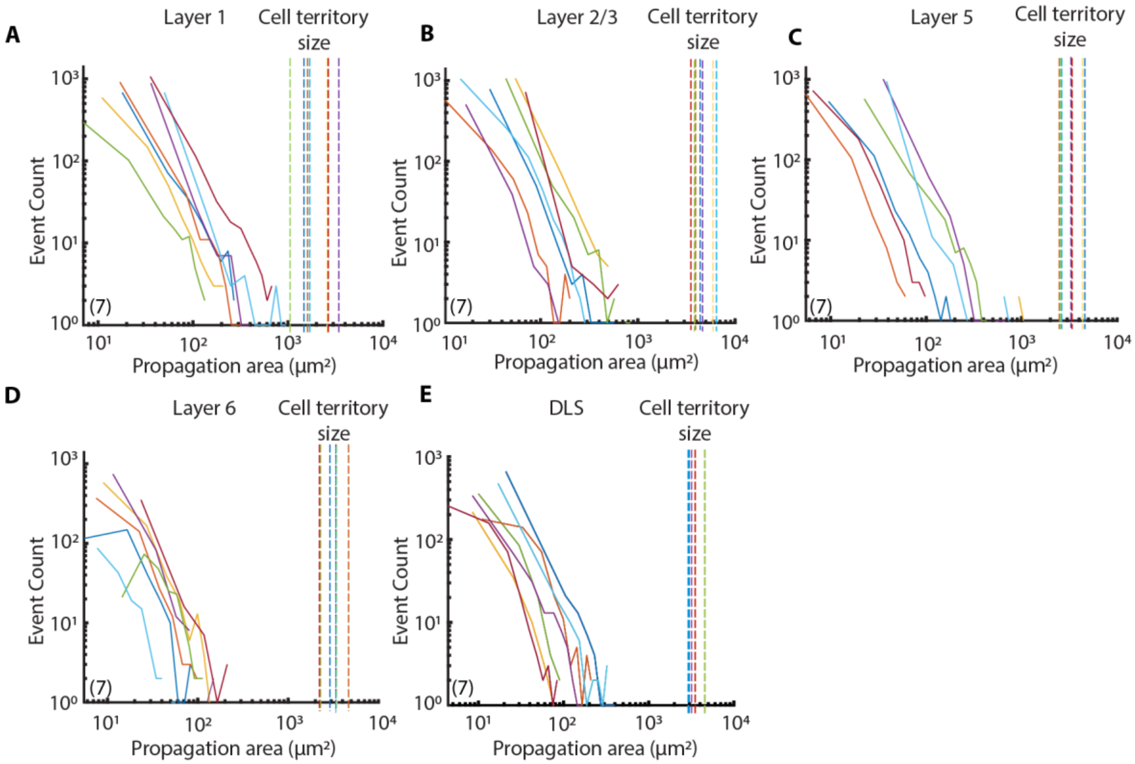


**Figure S4. Region-dependent variation in event count with respect to spreading size.**

(A-E) Plot showing the event count plotted against their spreading/propagation size. Each colored dotted line represents the respective territory size of the recorded astrocytes reported in this plot. (n = 7 (L1), n = 7 (L2/3), n = 7 (L5), n = 7 (L6), n = 7 (DLS) astrocytes; N = 14 mice).

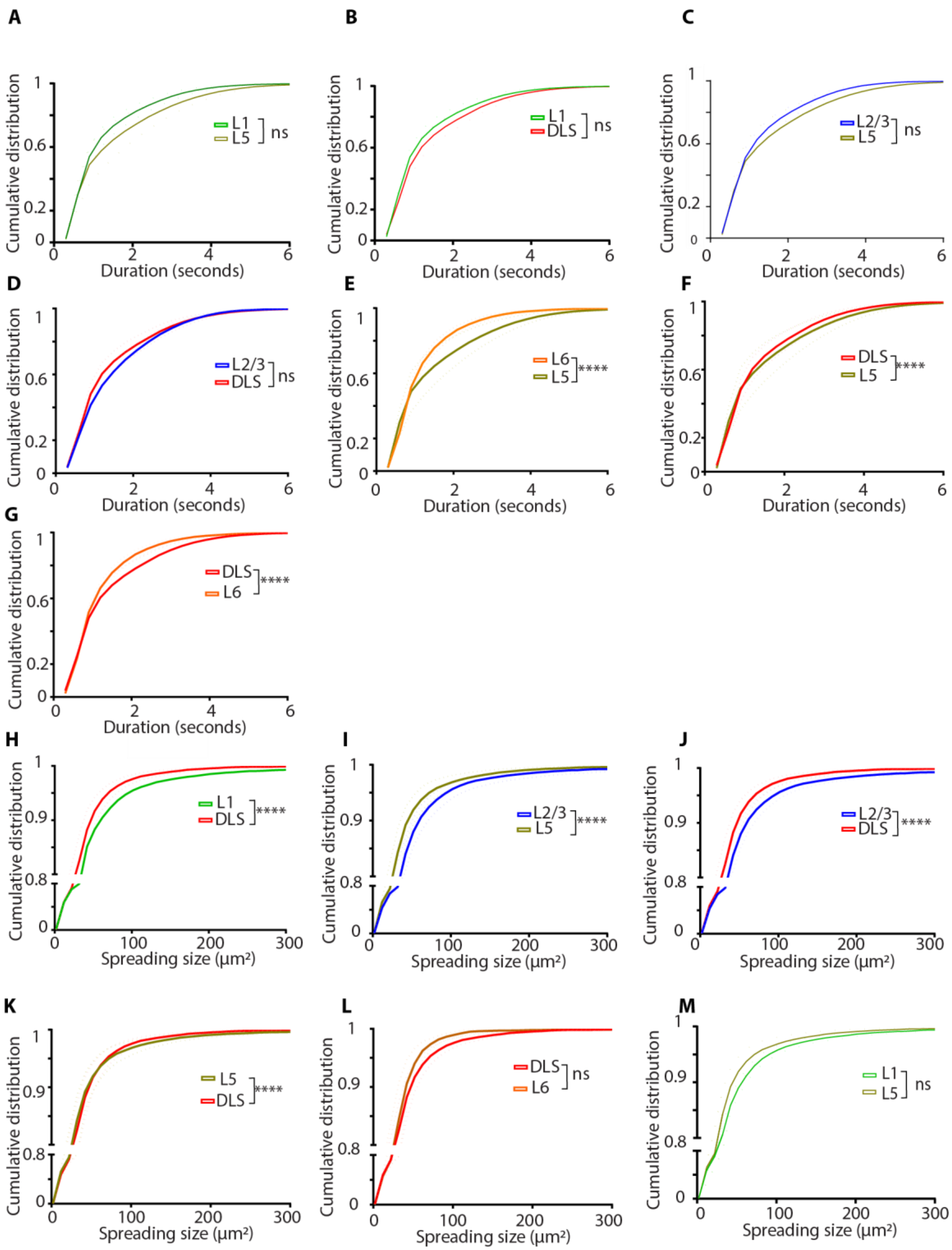


**Figure S5. Pairwise cumulative distributions of event duration and spreading size across the MOp and the DLS.**

(A-G) Pairwise comparison of CDF of event duration. (n = 12 (L1), n = 11 (L2/3), n = 7 (L5), n = 7 (L6), n = 7 (DLS) astrocytes; N = 19 mice; Kolmogorov-Smirnov's test).
(H-M) Pairwise comparison of CDF of event spreading/propagation size. (n = 12 (L1), n = 11 (L2/3), n = 7 (L5), n = 7 (L6), n = 7 (DLS) astrocytes; N = 19 mice; Kolmogorov-Smirnov's test).

The data are shown as mean ± SEM.
****$p < 0.0001$; ns = non-significant.

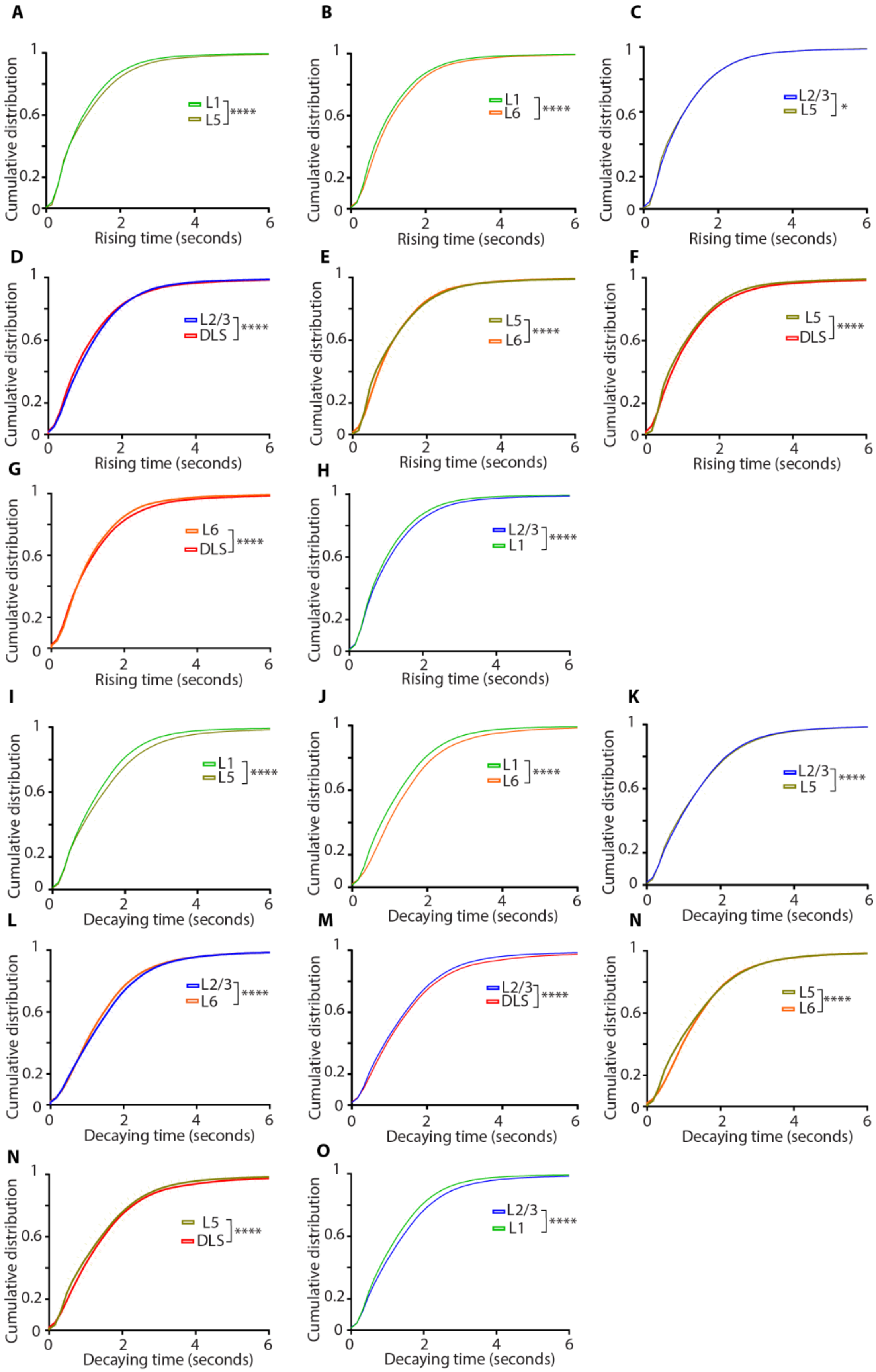
A
Cumulative distribution
Rising time (seconds)
L1
L5
****
B
Cumulative distribution
Rising time (seconds)
L1
L6
****
C
Cumulative distribution
Rising time (seconds)
L2/3
L5
*
D
Cumulative distribution
Rising time (seconds)
L2/3
DLS
****
E
Cumulative distribution
Rising time (seconds)
L5
L6
****
F
Cumulative distribution
Rising time (seconds)
L5
DLS
****
G
Cumulative distribution
Rising time (seconds)
L6
DLS
****
H
Cumulative distribution
Rising time (seconds)
L2/3
L1
****
I
Cumulative distribution
Decaying time (seconds)
L1
L5
****
J
Cumulative distribution
Decaying time (seconds)
L1
L6
****
K
Cumulative distribution
Decaying time (seconds)
L2/3
L5
****
L
Cumulative distribution
Decaying time (seconds)
L2/3
L6
****
M
Cumulative distribution
Decaying time (seconds)
L2/3
DLS
****
N
Cumulative distribution
Decaying time (seconds)
L5
L6
****
N
Cumulative distribution
Decaying time (seconds)
L5
DLS
****
O
Cumulative distribution
Decaying time (seconds)
L2/3
L1
****

**Figure S6. Pairwise cumulative distributions of event rising and decay time across the MOp and the DLS.**

(A-H) Pairwise comparison of CDF of rising time. (n = 12 (L1), n = 11 (L2/3), n = 7 (L5), n = 7 (L6), n = 7 (DLS) astrocytes; N = 19 mice; Kolmogorov-Smirnov's test).
(I-O) Pairwise comparison of CDF of event decaying time. (n = 12 (L1), n = 11 (L2/3), n = 7 (L5), n = 7 (L6), n = 7 (DLS) astrocytes; N = 19 mice; Kolmogorov-Smirnov's test).
The data are shown as mean ± SEM.
*$p < 0.05$, ****$p < 0.0001$.

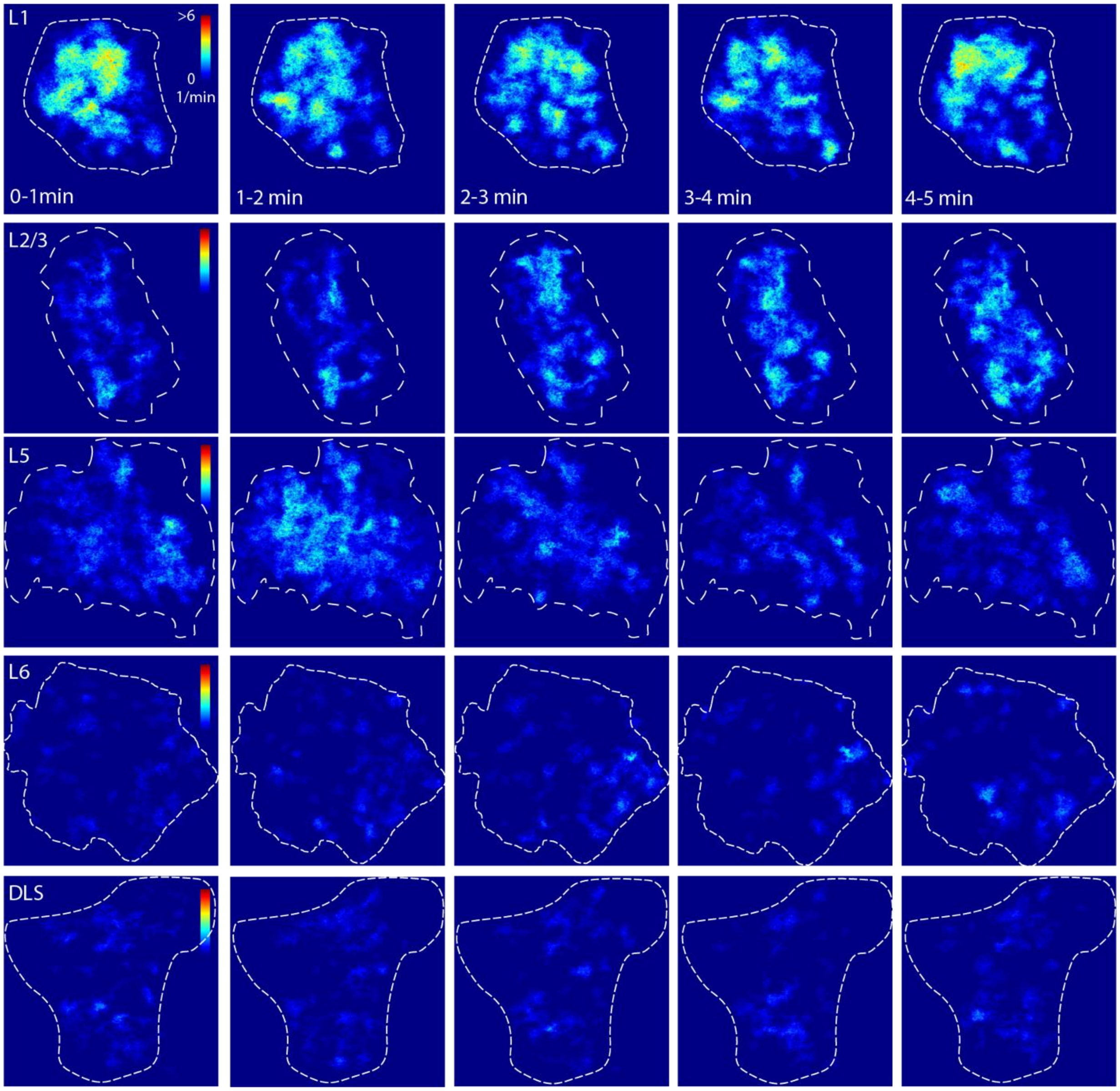


**Figure S7. Hotspot pattern exhibition during every minute of recording of astrocytes across all regions.**

Each row represents a single astrocyte recorded from L1, L2/3, L5, L6 and DLS. Each column represents the hotspot pattern in every minute across 5 minutes of recording. The white dashed line marks the territory of the astrocyte. Scale bar represents 20 μm.

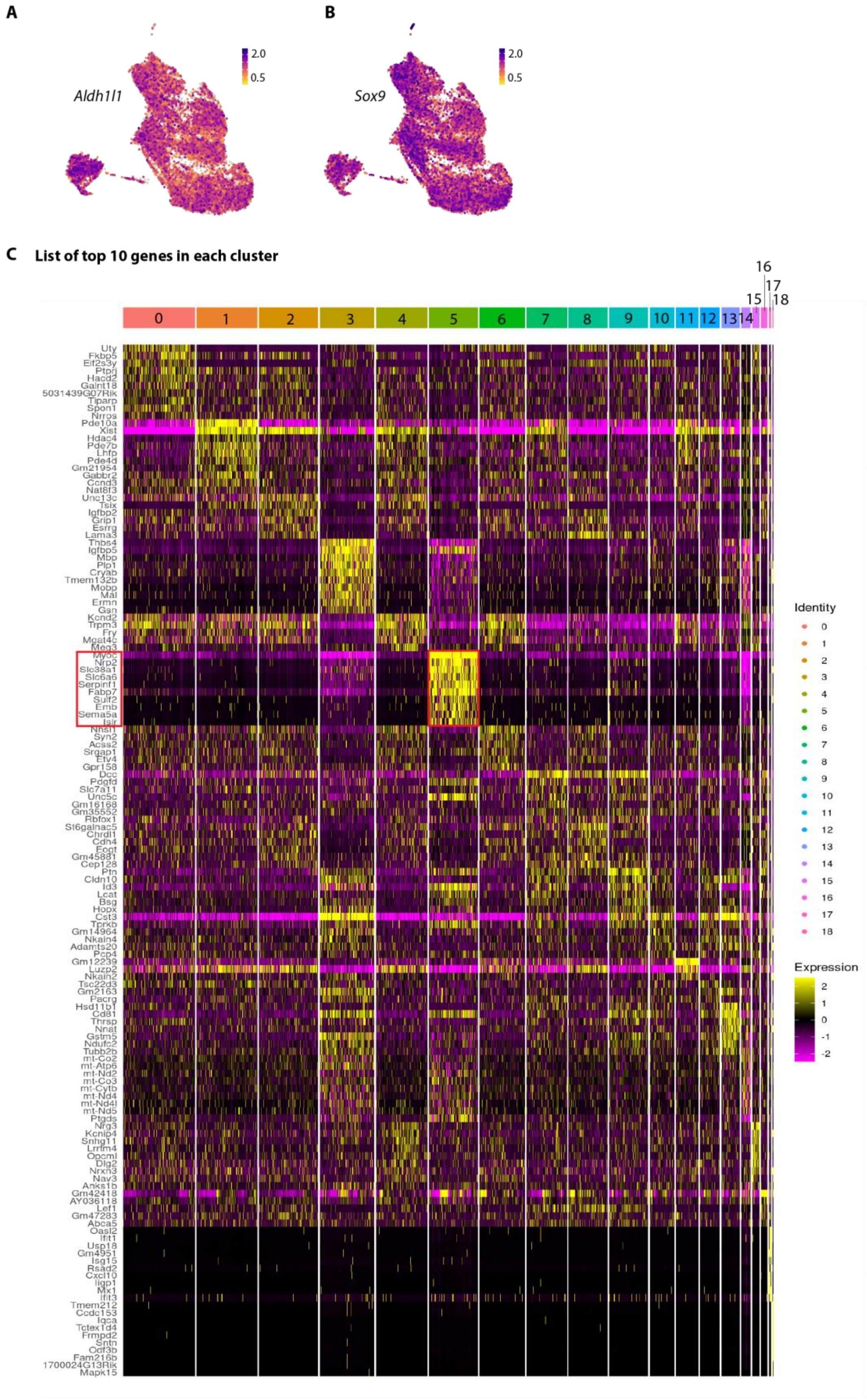
A
Aldh1l1
2.0
0.5
B
Sox9
2.0
0.5
C
List of top 10 genes in each cluster
Identity
Expression

**Figure S8. List of the top 10 expressed genes in each cluster of the astrocyte dataset.**

(A) UMAP showing the expression of *Aldh1l1* gene in the astrocyte cluster.
(B) UMAP showing the expression of *Sox9* gene in the astrocyte cluster.
(C) List of top 10 genes in each cluster. The red box highlights the top 10 genes in cluster 5.

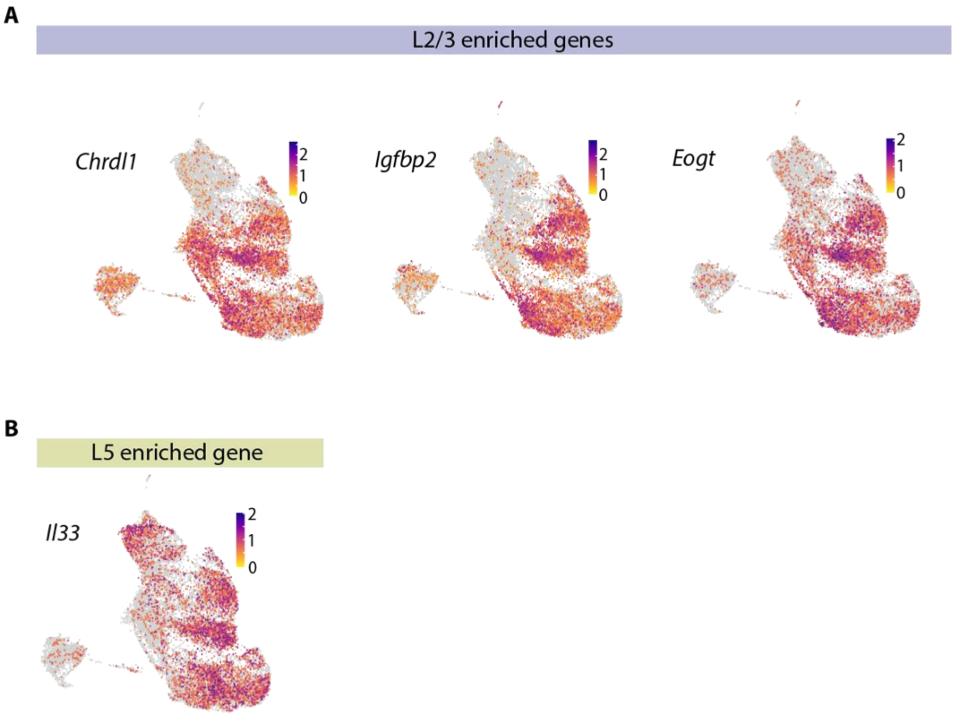


**Figure S9. List of L2/3 and L5 astrocyte enriched genes.**

(A) MOp astrocytic expression pattern of genes reported to show enriched expression in L2/3 astrocytes within the SSp[3].

(B) MOp astrocytic expression pattern of the gene reported to show enriched expression in L5 astrocytes within the SSp[3].

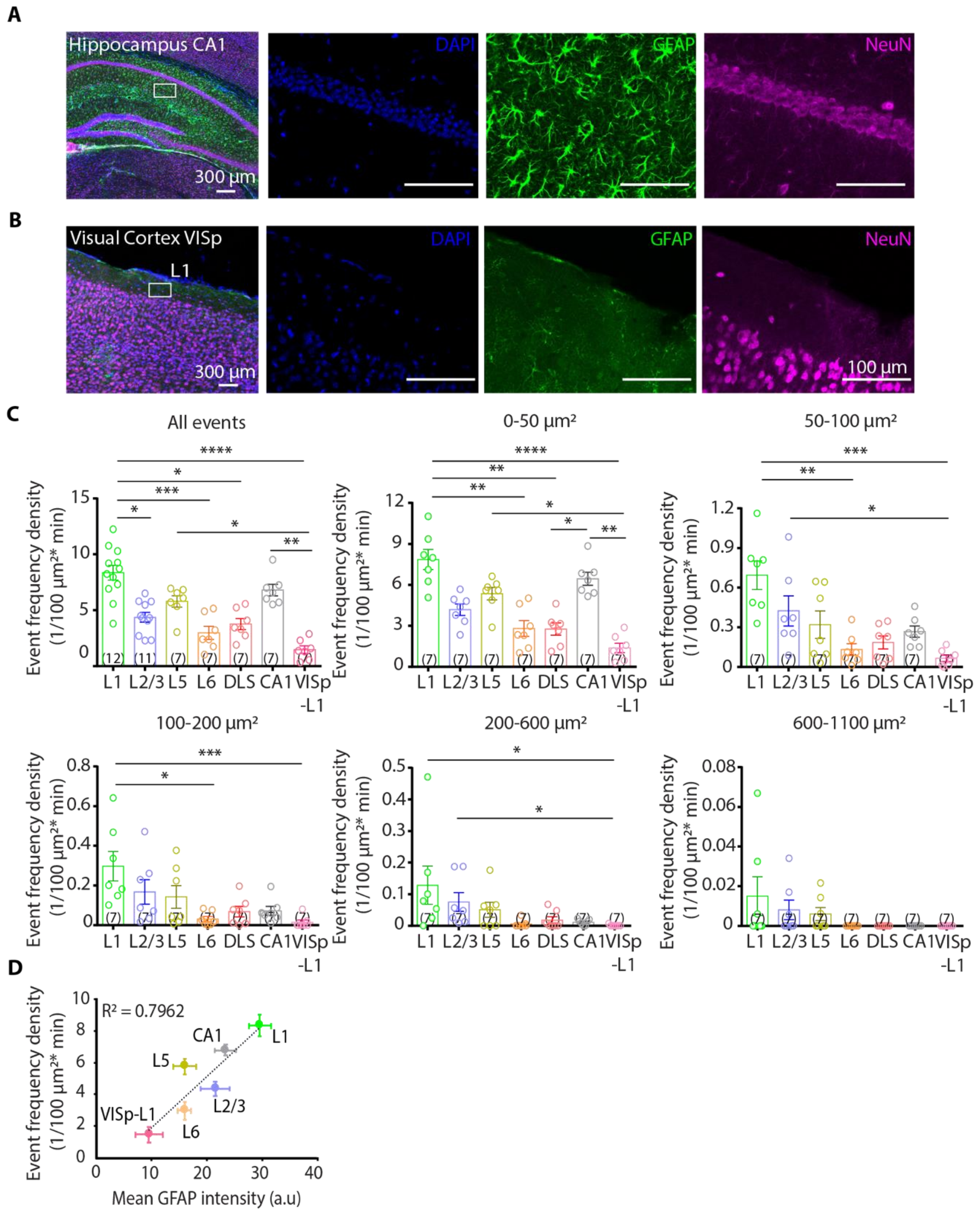


**Figure S10. Event frequency density across spreading size subgroups in different brain regions.**

(A) Histology studies showing the GFAP (green), NeuN (magenta) and DAPI (blue) staining the CA1 region.
(B) Histology studies showing the GFAP (green), NeuN (magenta) and DAPI (blue) staining the V1-L1 region. Scale represents 300µm and 100 µm.

(C) Event frequency density of spreading subgroups, quantified across the MOp, DLS, CA1 and the VISp-L1. (n = 12 (L1), n = 11 (L2/3), n = 7 (L5), n = 7 (L6), n = 7 (DLS), n = 7 (CA1), n = 7 (VISp-L1) astrocytes; N = 24 mice, Kruskal-Wallis test followed by Dunn’s multiple comparisons test).

(All events (Unit: events/$10^2$ $\mu m^2$ per minute)**;** L1: 0.0835 ± 0.0066, n = 12, L2/3: 0.0436 ± 0.0043, n = 11, L5: 0.0579 ± 0.0050, n = 7, L6: 0.0298 ± 0.0058, n = 7, DLS: 0.0375 ± 0.0049, n = 7, CA1: 0.0680 ± 0.0051, n = 7, VISp-L1: 0.0147 ± 0.0034, n = 7; 0-50 $\mu m^2$ (Unit: events/$10^2$ $\mu m^2$ per minute)**;** L1: 0.07856 ± 0.0074, n = 7, L2/3: 0.0418 ± 0.0042, n = 7, L5: 0.0535 ± 0.0045, n = 7, L6: 0.0281 ± 0.0057, n = 7, DLS: 0.0276 ± 0.0044, n = 7, CA1: 0.0645 ± 0.0047, n = 7, VISp-L1: 0.0139 ± 0.0034, n = 7; 50-100 $\mu m^2$ (Unit: events/$10^2$ $\mu m^2$ per minute); L1: 0.0069 ± 0.0010, n = 7, L2/3: 0.0042 ± 0.0011, n = 7, L5: 0.0032 ± 0.0010, n = 7, L6: 0.0013 ± 0.0004, n = 7, DLS: 0.0018 ± 4 x $10^{-4}$, n = 7, CA1: 0.0026 ± 4 x $10^{-4}$, n = 7, VISp-L1: 6 x $10^{-4}$ ± 2 x $10^{-4}$, n = 7; 100-200 $\mu m^2$ (Unit: events/$10^2$ $\mu m^2$ per minute); L1: 0.0029 ± 7 x $10^{-4}$, n = 7, L2/3: 0.0016 ± 6 x $10^{-4}$, n = 7, L5: 0.0014 ± 5 x $10^{-4}$, n = 7, L6: 3 x $10^{-4}$ ± 1 x $10^{-4}$, n = 7, DLS: 6 x $10^{-4}$ ± 2 x $10^{-4}$, n = 7, CA1: 6 x $10^{-4}$ ± 2 x $10^{-4}$, n = 7, VISp-L1: 1 x $10^{-4}$ ± 1 x $10^{-4}$, n = 7; 200-600 $\mu m^2$ (Unit: events/$10^2$ $\mu m^2$ per minute); L1: 0.0012 ± 6 x $10^{-4}$, n = 7, L2/3: 7 x $10^{-4}$ ± 2 x $10^{-4}$, n = 7, L5: 5 x $10^{-4}$ ± 2 x $10^{-4}$, n = 7, L6: 5 x $10^{-5}$ ± 4 x $10^{-5}$, n = 7, DLS: 1 x $10^{-4}$ ± 1 x $10^{-4}$, n = 7, CA1: 1 x $10^{-4}$ ± 3 x $10^{-5}$, n = 7, VISp-L1: 1 x $10^{-5}$ ± 1 x $10^{-5}$, n = 7; 600-1100 $\mu m^2$ (Unit: events/$10^2$ $\mu m^2$ per minute); L1: 1 x $10^{-4}$ ± 9 x $10^{-5}$, n = 7, L2/3: 8 x $10^{-5}$ ± 4 x $10^{-5}$, n = 7, L5: 6 x $10^{-5}$ ± 3 x $10^{-5}$, n = 7, L6: 0 ± 0, n = 7, DLS: 0 ± 0, n = 7, CA1: 0 ± 0, n = 7, VISp-L1: 0 ± 0, n = 7 Kruskal-Wallis test followed by Dunn’s multiple comparisons test).

(D) Correlation between mean GFAP density (a.u) with their event frequency density. (Y-axis: n = 7 astrocytes each; N = 19 mice, X-axis: n = 6 slices; N = 3 mice).

The data are shown as mean ± SEM.

*p < 0.05, **p < 0.01, ***p < 0.001, ****p < 0.0001.

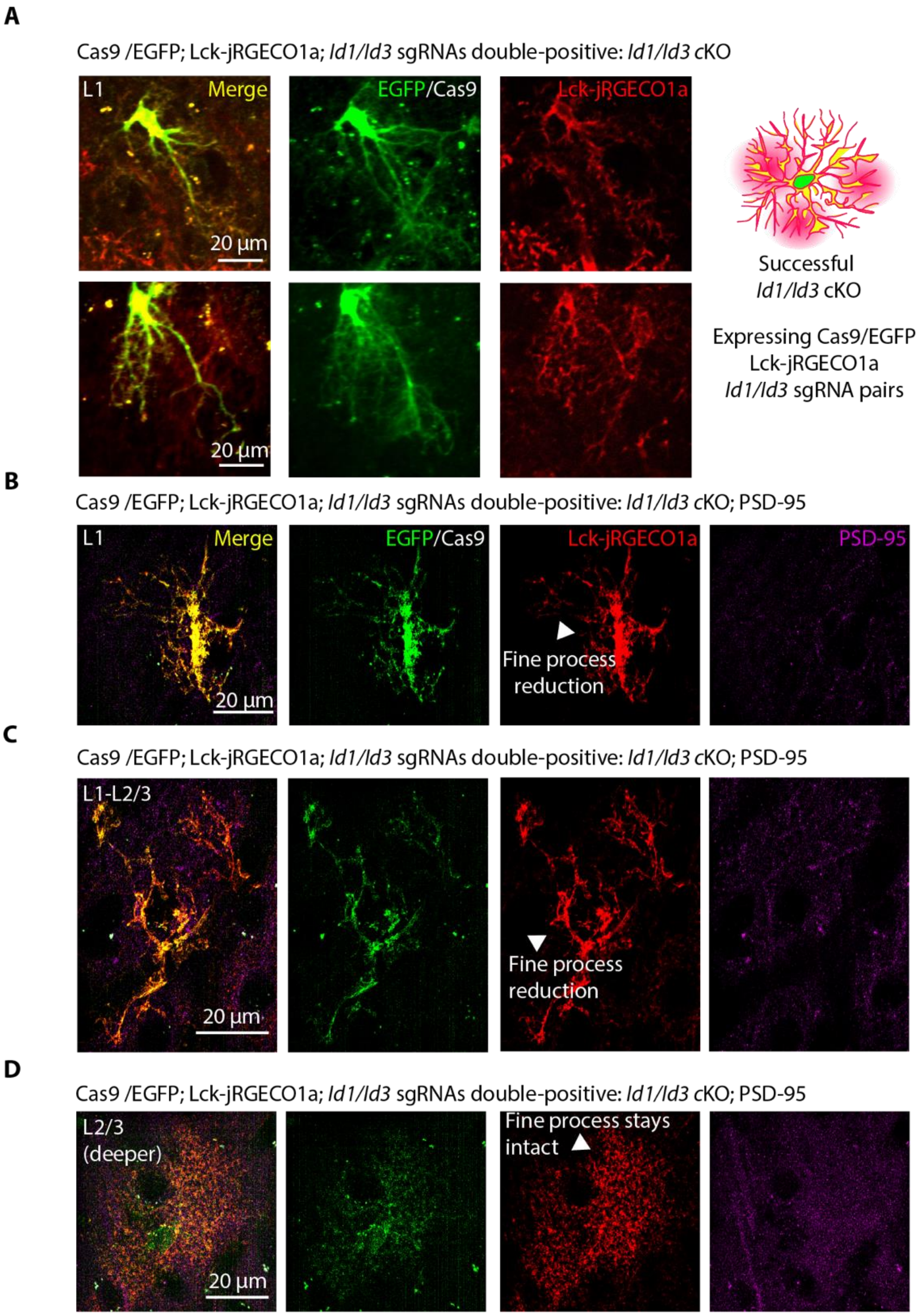


**Figure S11. *Id1/Id3* genes cKO reduces fine processes and territorial complexity in L1-L2/3 astrocytes.**

(A) (Left) Examples of L1 astrocytes upon cKO of *Id1/Id3* genes, imaged using two-photon microscopy. EGFP/Cas9 (green), Lck-jRGECO1a (red). (Right) Illustration showing an astrocyte with successful cKO, expressing both Lck-jRGECO1a and EGFP/Cas9.
(B) L1 astrocyte upon cKO of *Id1/Id3* genes, imaged using SR-SIM. EGFP/Cas9 (green), Lck-jRGECO1a (red).

(C) L1-L2/3 transition residing astrocyte upon cKO of *Id1/Id3* genes, imaged using SR-SIM. EGFP/Cas9 (green), Lck-jRGECO1a (red), PSD-95 (magenta).

(D) Deeper residing L2/3 astrocyte upon cKO of *Id1/Id3* genes, imaged using SR-SIM. EGFP/Cas9 (green), Lck-jRGECO1a (red), PSD-95 (magenta).
